# Non-lesional and Lesional Lupus Skin Share Inflammatory Phenotypes that Drive Activation of CD16+ Dendritic Cells

**DOI:** 10.1101/2021.09.17.460124

**Authors:** Allison C. Billi, Feiyang Ma, Olesya Plazyo, Mehrnaz Gharaee-Kermani, Rachael Wasikowski, Grace A. Hile, Xianying Xing, Christine M. Yee, Syed M. Rizvi, Mitra P. Maz, Fei Wen, Lam C. Tsoi, Matteo Pellegrini, Robert L. Modlin, Johann E. Gudjonsson, J. Michelle Kahlenberg

## Abstract

Cutaneous lupus erythematosus (CLE) is a disfiguring and poorly understood condition frequently associated with systemic lupus. Studies to date suggest that non-lesional keratinocytes play a role in disease predisposition, but this has not been investigated in a comprehensive manner or in the context of other cell populations. To investigate CLE immunopathogenesis, normal-appearing skin, lesional skin, and circulating immune cells from lupus patients were analyzed via integrated single-cell RNA-sequencing and spatial-seq. We demonstrate that normal-appearing skin of lupus patients represents a type I interferon-rich, ‘prelesional’ environment that skews gene transcription in all major skin cell types and dramatically distorts cell-cell communication. Further, we show that lupus-enriched CD16+ dendritic cells undergo robust interferon education in the skin, thereby gaining pro-inflammatory phenotypes. Together, our data provide a comprehensive characterization of lesional and non-lesional skin in lupus and identify a role for skin education of CD16+ dendritic cells in CLE pathogenesis.

## INTRODUCTION

Cutaneous lupus erythematosus (CLE) is a disfiguring inflammatory skin disease that affects 70% of patients with systemic lupus erythematosus (SLE). While about 50% of patients respond to SLE-directed therapies^1^, many patients suffer from refractory skin lesions, even when their systemic disease is controlled. Lack of knowledge regarding the inflammatory composition of CLE and the drivers that instigate disease has delayed effective therapy development.

Intriguingly, the etiology of skin lesions in CLE may be, at least partially, found in abnormalities in non-lesional, normal-appearing skin. Recent data support a role for increased epidermal type I interferon (IFN) production^2, 3, 4, 5^ and dysfunction of Langerhans cells^6^ as important for priming inflammatory and apoptotic responses. However, the role of other cells in the skin, the skewed communication networks between them, and cellular mediators of this inflammatory predisposition have not been well-defined.

In this paper, we examined the cellular composition of paired lesional and non-lesional skin samples from SLE patients with active CLE lesions to comprehensively define the cellular makeup and to characterize the principal mediators of inflammatory changes that contribute to the disease. We further examined the peripheral blood of the same patients to investigate the cutaneous education of monocyte-derived dendritic cells, which were found to be prominent in lesional and non-lesional skin. Overall, we found an IFN-rich signature and a unique, pro-inflammatory cellular communication network between stromal and inflammatory cells in lesional and non-lesional skin that supports a critical role for the skin itself in priming inflammatory responses in SLE patients.

## RESULTS

### Single-cell RNA-sequencing (scRNA-seq) of lesional and non-lesional skin from patients with CLE identifies diverse skin and immune cell populations

To investigate the cellular composition and comprehensive transcriptional effects of CLE, we performed scRNA-seq on lesional and sun-protected non-lesional skin from 7 patients with active CLE (**Supplementary Table 1**), 6 of whom also carried a diagnosis of SLE. Samples were analyzed in parallel with skin from 14 healthy controls from diverse sites. The final dataset comprised 46,540 cells, with an average of 2,618 genes and 11,645 transcripts per cell. Visualization using Uniform Manifold Approximation and Projection (UMAP) revealed 26 distinct cell clusters (**Fig. 1a**) that were annotated as 10 major cell types, each comprising cells from lesional, non-lesional, and healthy control skin biopsies (**Fig. 1b-d**). Conspicuous clustering by disease state was evident for many cell types, including keratinocytes (KCs), myeloid cells, and melanocytes. Cell composition analysis revealed an increase in the proportion of myeloid cells in both lesional and non-lesional skin relative to healthy control (**Fig. 1e**).

**Figure 1.**
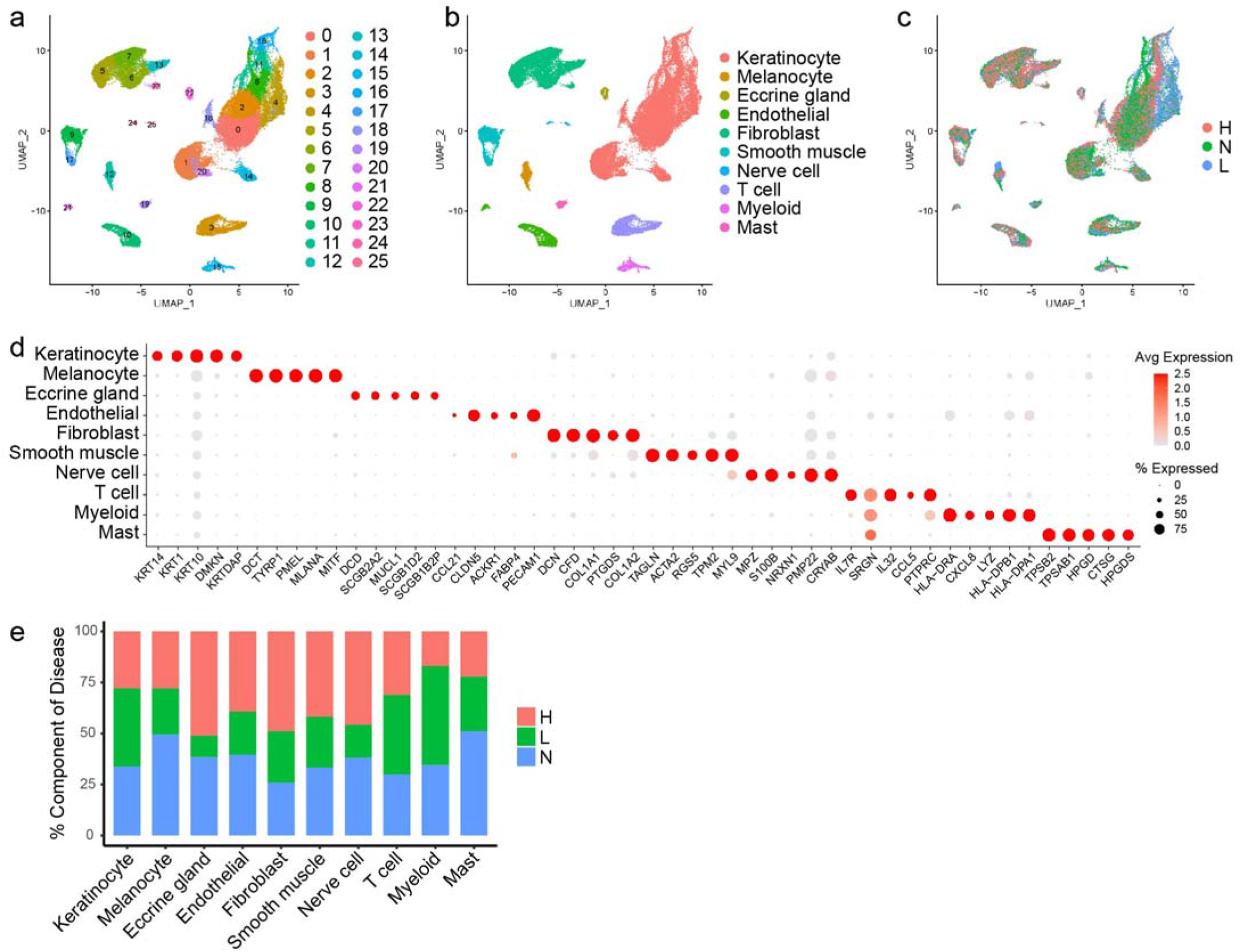
Single-cell RNA-sequencing (scRNA-seq) captures the cellular diversity within lesional and non-lesional skin of patients with cutaneous lupus erythematosus (CLE). **a.** UMAP plot of 46,540 cells colored by cluster. **b.** UMAP plot of cells colored by cell type. **c.** UMAP plot of cells colored by disease state. H, healthy control skin. N, non-lesional lupus skin. L, lesional lupus skin. **d.** Dot plot of representative marker genes for each cell type. Color scale, average marker gene expression. Dot size, percentage of cells expressing marker gene. **e.** Bar plot of cell type proportions across disease states.

### KCs from both lesional and non-lesional skin of patients with CLE exhibit a pathologic type I IFN signature

KCs constituted the majority of cells sequenced (25,675 cells). Sub-clustering analysis of KCs identified 14 sub-clusters (**Fig. 2a**), including several (5, 6, 8, 13) dominated by KCs from lupus patients (**Fig. 2b,c**). Analyses of characteristic KC subtype markers identified 5 KC states: basal, spinous, supraspinous, follicular, and cycling (**Fig. 2d, Supplementary Fig. 1a**). Lupus-dominated sub-clusters corresponded to subpopulations within basal and spinous KC states.

**Figure 2.**
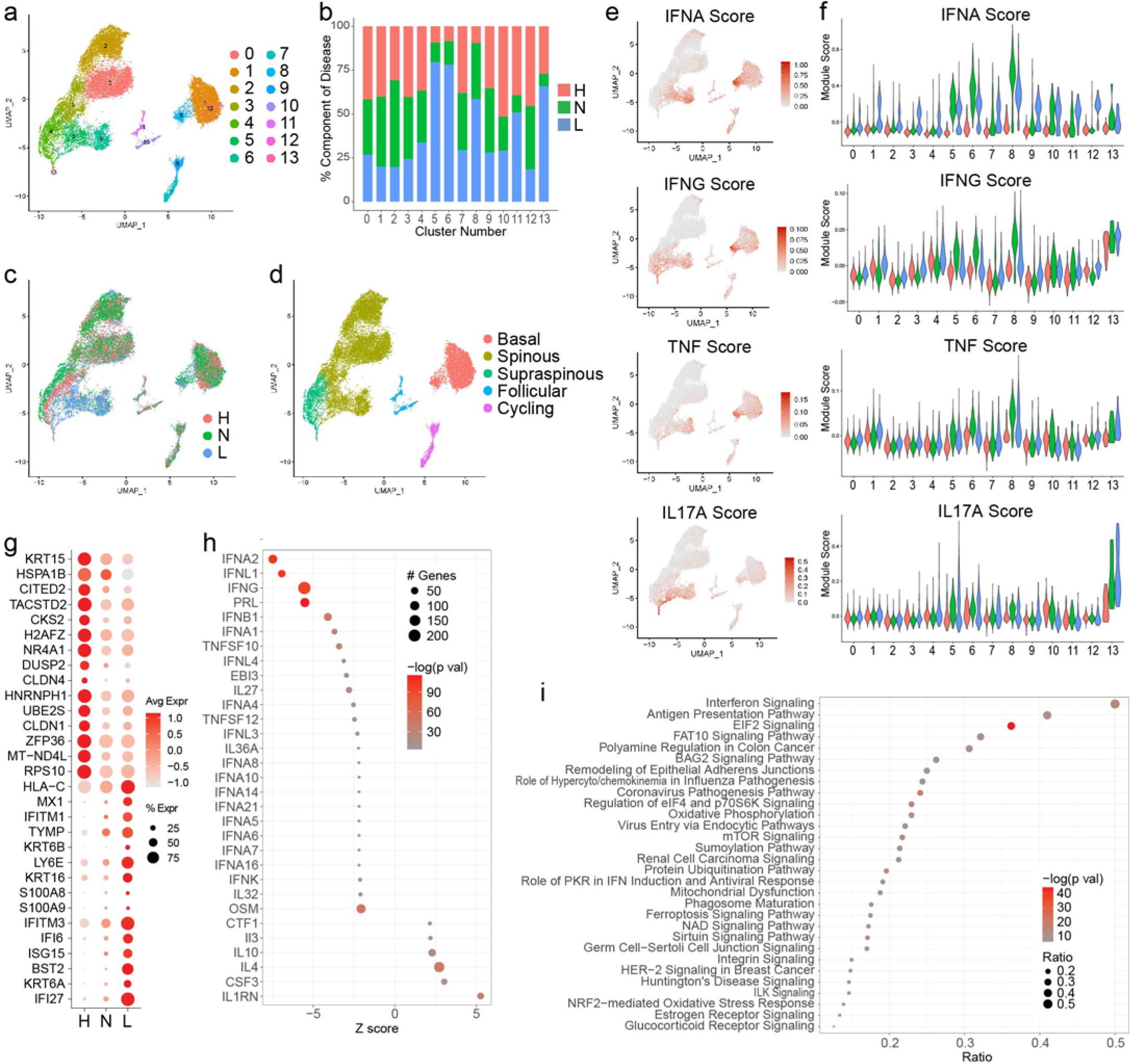
Interferon (IFN) responses shape the transcriptomic landscape of both lesional and non-lesional keratinocytes in patients with CLE. **a.** UMAP plot of 25,675 keratinocytes (KCs) colored by sub-cluster. **b.** Bar plot of number of KCs in each sub-cluster split by disease state. **c.** UMAP plot of KCs colored by disease state. **d.** UMAP plot of KCs colored by subtype. **e.** Feature plots of module scores for the indicated KC cytokine modules. **f.** Violin plots of KC scores for the indicated cytokine modules split by disease state. **g.** Dot plot of the top 15 differentially expressed genes (DEGs) downregulated and upregulated in L vs. H basal KCs. Color scale, average marker gene expression. Dot size, percentage of cells expressing marker gene. **h.** Dot plot of the top 30 upstream regulators enriched among DEGs in L vs. H basal KCs. Color scale, −log10(p value) from the enrichment analysis. Dot size, number of DEGs corresponding to each upstream regulator. Negative Z score, enriched in L; positive, enriched in H. **i.** Dot plot of the top 30 canonical pathways enriched among DEGs in L vs. H basal KCs. Color scale, −log10(p value) from the enrichment analysis. Dot size, ratio of [number of pathway genes among the DEGs]/[total number of pathway genes].

The relatively shallow depth of scRNA-seq precludes direct examination of transcript levels for many cytokines implicated in CLE – particularly IFNs. Thus, to investigate whether cytokine responses were driving this KC sub-clustering by disease, we calculated KC module scores derived from genes induced in cultured KCs upon stimulation with the indicated cytokines and generated feature plots displaying module scores for each cytokine (**Fig. 2e, Supplementary Fig. 1b**). Lupus-enriched sub-clusters corresponded best with cells exhibiting high scores for type I IFN (IFN-α), type II IFN (IFN-γ), and to a lesser degree TNF. Our and others’ prior work has identified a critical role for type I IFN in SLE and CLE keratinocytes^2, 3, 4, 5^. Accordingly, cytokine module violin plots revealed that lupus-enriched basal (sub-cluster 8) and spinous (5 and 6) sub-clusters consisted almost entirely of KCs with high IFN-α module scores (**Fig. 2f, Supplementary Fig. 1c**), whereas the separation was less striking for IFN-γ and TNF. Notably, cells scoring highest in these clusters originated from non-lesional biopsies. This suggests that even normal-appearing skin from patients with CLE exists in a ‘prelesional’ state, primed by heightened type I IFN signaling. Follicular KC sub-clusters (10 and 11) showed elevated IFN-α cytokine module scores in non-lesional and lesional samples relative to healthy control as well (**Fig. 2f**), suggesting the follicular epithelium also represents an abnormal, IFN-rich environment in CLE; however, scores were far lower for follicular than basal KCs, implicating the interfollicular epidermis more strongly than the follicular epithelium in type I IFN education of neighboring stromal and skin-infiltrating cells.

For a broader understanding of the transcriptomic differences in lesional KCs of patients with CLE, we performed differential expression analysis between the non-lesional and lesional CLE vs. healthy basal KCs and identified type I IFN downstream genes (e.g., *MX1*, *IFITM1*, *IFITM3*, *IFI6*, *ISG15*, *IFI27*) among the top upregulated genes in the lesional cells (**Fig. 2g**). We then used Ingenuity Pathway Analysis (IPA) to identify the top cytokines predicted to serve as upstream regulators for the genes induced in lesional samples, identifying primarily IFNs as upstream regulators of CLE-enriched transcripts (**Fig. 2h**). Corroborating this, canonical pathway analysis distinguished IFN signaling as highly enriched in lesional samples (**Fig. 2i**).

### scRNA-seq identifies a CLE-enriched fibroblast subtype exhibiting a strong IFN response signature and IL-17A influence restricted to lesional skin

We next analyzed fibroblasts (FBs), the other major stromal cell constituent of the skin. Sub-clustering analysis of 8,622 FBs identified 10 sub-clusters (**Fig. 3a**). Only one sub-cluster (4) was dominated by FBs from lupus patients (**Fig. 3b,c**). Annotation of these sub-clusters based on published dermal FB marker genes^7^ revealed three subtypes as previously described (SFRP2+, COL11A1+, and SFRP4+ FBs) and a small cluster marked by expression of *COL66A1* and *RAMP1* (RAMP1+ FBs) (**Fig. 3d, Supplementary Fig. 2a**). Immunohistochemistry of these key markers (**Supplementary Fig. 2b**) confirmed that SFRP2+ FBs constituted the majority of FBs^7^. The lupus-enriched sub-cluster lay within the SFRP2+ FBs and was analyzed as an independent subtype. Analysis of the top gene markers of each FB subtype indicated that these FBs were distinguished by high IFN-stimulated gene (ISG) expression (**Supplementary Fig. 2c**), and thus we designated these IFN FBs. Consistent with this, feature and violin plots depicting FB cytokine module scores calculated using genes induced in cultured FBs stimulated by cytokines as above revealed that IFN FBs were most uniquely distinguished by IFN-α and IFN-γ cytokine signatures (**Fig. 3e,f, Supplementary Fig. 2d,e**). As in KCs (**Fig. 2e**), IFN cytokine module scores were highest in non-lesional FBs, reinforcing that normal-appearing skin of lupus patients represents a prelesional, IFN-primed environment. We compared the cytokine upstream regulators identified in the comparisons of non-lesional vs. healthy basal KCs and non-lesional IFN FBs vs. healthy SFRP2+ FBs. This revealed that type I IFNs, such as IFNA2, IFNL1, and IFNB1, served as the top upstream regulators in both cell types (**Fig. 3g**). Notably, high FB IFN module scores were primarily restricted to the single sub-cluster of IFN FBs (**Fig. 3f**), indicating that only a specific subset of FBs in skin of patients with CLE exhibits robust IFN education. This is an interesting contrast to KCs, where non-lesional and/or lesional KCs showed elevated IFN module scores for the majority of sub-clusters (**Fig. 2e**).

**Figure 3.**
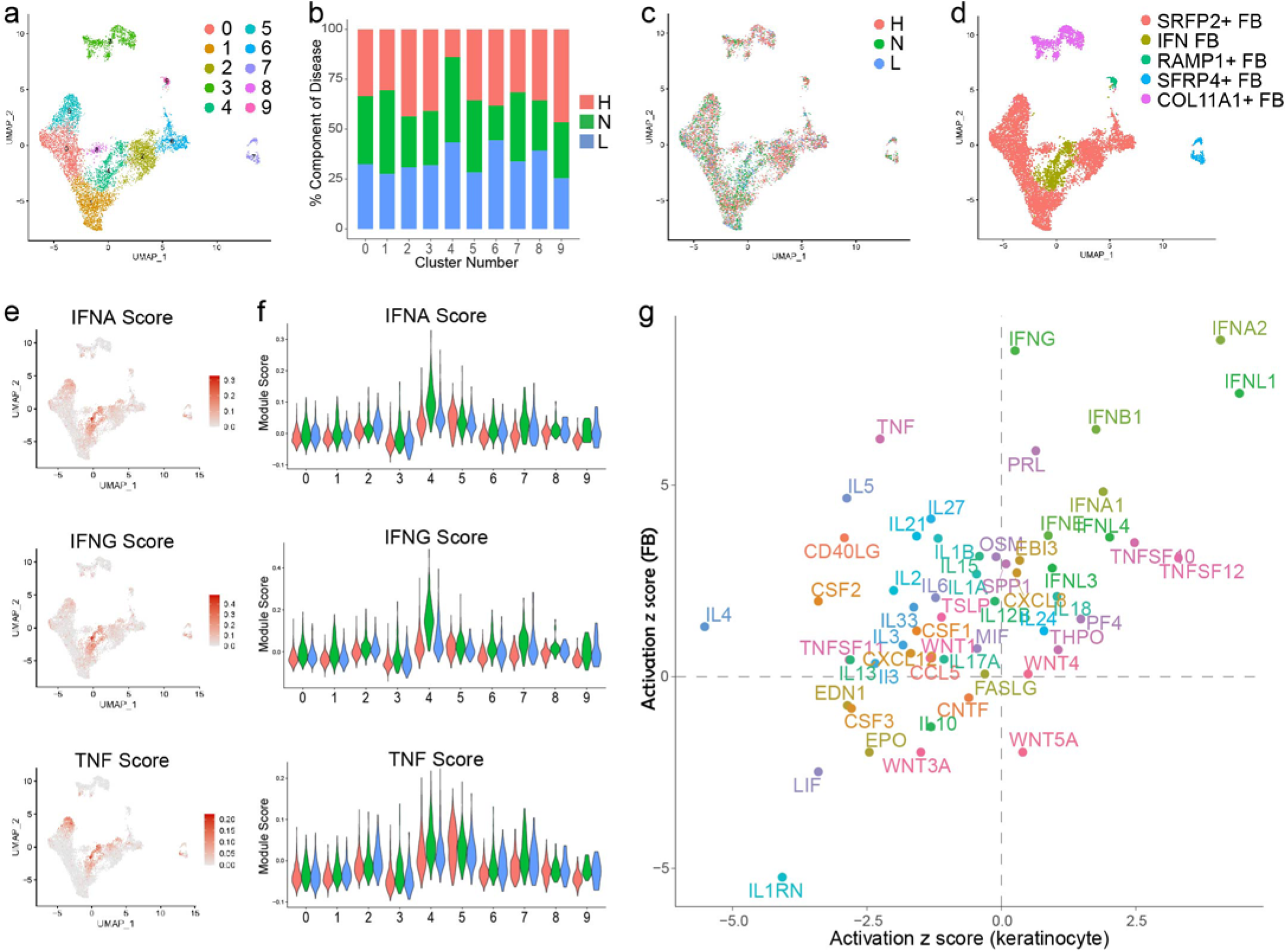
Analysis of fibroblast (FB) heterogeneity in CLE patient skin identifies an IFN-educated FB subtype present in both non-lesional and lesional skin. **a.** UMAP plot of 8,622 FBs colored by sub-cluster. **b.** Bar plot of number of FBs in each sub-cluster split by disease state. **c.** UMAP plot of FBs colored by disease state. **d.** UMAP plot of FBs colored by subtype. **e.** Feature plots of scores for the indicated FB cytokine module scores. **f.** Violin plots of FB module scores for the indicated cytokine modules split by disease state. **g.** Scatter plot showing activation z scores for cytokine upstream regulators common to basal KCs (N vs. H) and IFN FBs (N) vs. SFRP2+ FBs (H).

### T cells infiltrating lesional and non-lesional skin of patients with CLE demonstrate IFN education across multiple subsets including regulatory T cells

Having analyzed the major stromal cell types of the skin, we moved on to examination of the immune cells. Sub-clustering and annotation based on established marker genes identified nine T cell subsets (**Fig. 4a-c**). Cells of one sub-cluster were distinguished by expression of ISGs and were therefore designated IFN T cells (**Fig. 4c**). IFN T cells derived primarily from lupus samples (**Fig. 4d**), constituting 13% and 15% of T cells from non-lesional and lesional samples, respectively, but <1% of healthy control T cells.

**Figure 4.**
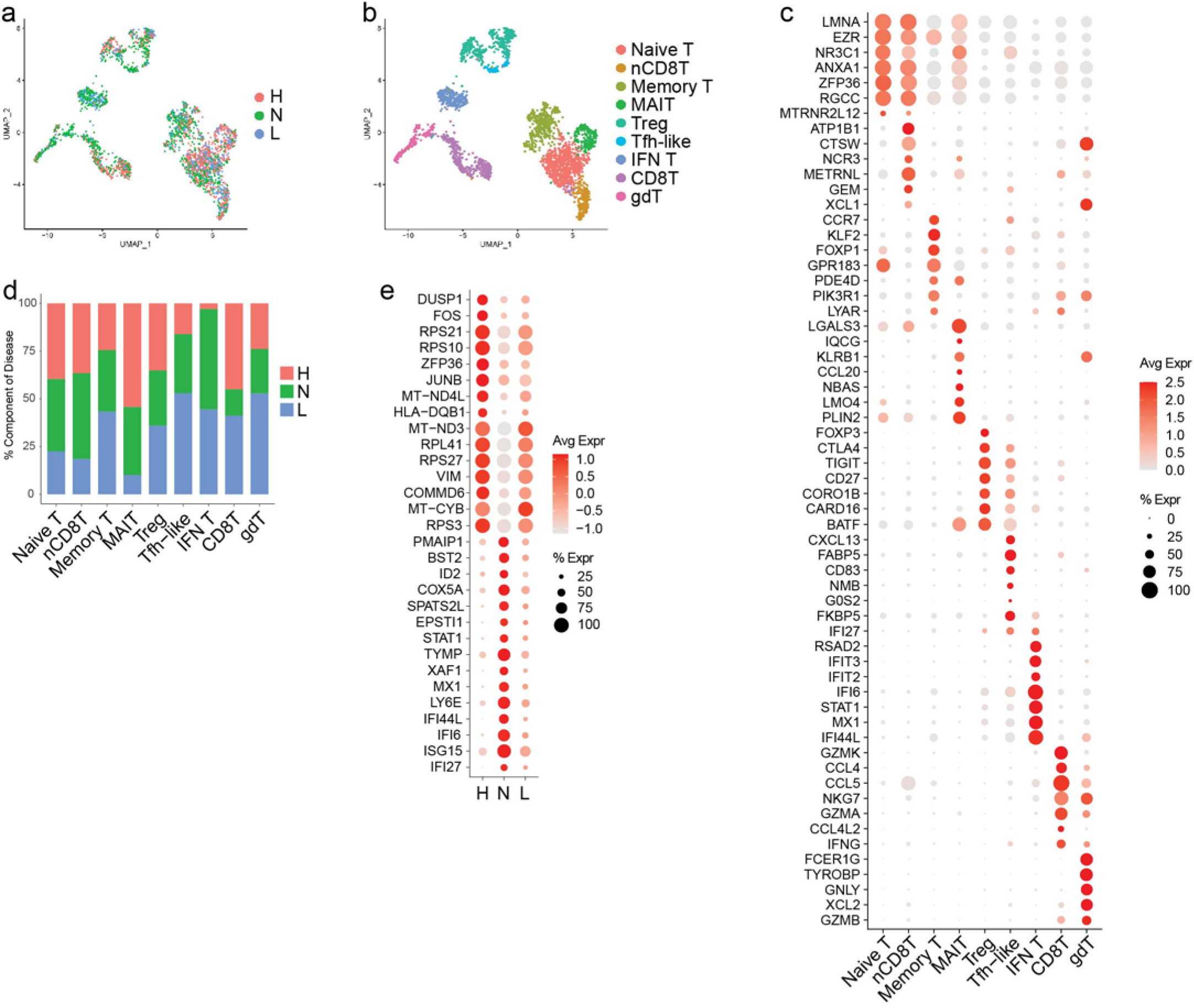
Skin of CLE patients exhibits an abnormal T cell infiltrate at both lesional and non-lesional sites. **a.** UMAP plot of 3,030 T cells colored by disease state. **b.** UMAP plot of T cells colored by subset. Naïve T, naïve CD4^+^ T cells; nCD8T, naïve CD8^+^ T cells; Memory T, memory T cells; MAIT, mucosal-associated invariant T cells; Treg, regulatory T cells; Tfh-like, T follicular helper-like cells; IFN T, interferon T cells; CD8T, CD8^+^ T cells; gdT, γδ T cells. **c.** Dot plot of representative marker genes for each T cell subset. Color scale, average marker gene expression. Dot size, percentage of cells expressing marker gene. **d.** Bar plot of disease state representation within each T cell subset. **e.** Dot plot of the top 15 differentially expressed genes (DEGs) downregulated and upregulated in N vs. H Tregs.

Regulatory T cells (Tregs), annotated based on *FOXP3* expression, were detected in similar proportions across healthy control, non-lesional, and lesional samples (**Fig. 4d**). Investigations of Treg abundance in peripheral blood of patients with SLE have yielded conflicting results^8, 9, 10, 11^, possibly due to differences in methodology or definition of Tregs^12^, but there appears to be consensus that Treg function is altered in patients with SLE^8, 9^. To investigate whether the non-lesional skin environment might influence Treg function in lupus patients, we examined the top DEGs upregulated in non-lesional vs. healthy control Tregs. This identified numerous ISGs, suggestive of chronic IFN stimulation (**Fig. 4e**). Overproduction of type I IFNs by antigen-producing cells (APCs) in SLE has been proposed as a cause of Treg dysfunction in SLE^13, 14^, and we have previously demonstrated that lupus-prone NZM2328 mice treated with ultraviolet light exhibit type I IFN-dependent suppression of Treg function^13^. Thus, the chronic IFN stimulation of Tregs that we observe in non-lesional skin may contribute to impaired Treg ability to maintain immune homeostasis and self-tolerance in lupus.

One sub-cluster of T cells clustered closely with Tregs but did not express *FOXP3*. Rather, this sub-cluster was distinguished by expression of *CXCL13* (**Fig. 4c**), a B cell-attracting chemokine and SLE biomarker that appears to play a pathogenic role (reviewed in *Schiffer et al., 2015*^15^), as well as *ICOS* and *PDCD1* (encoding PD-1), leading us to annotate these as T follicular helper (Tfh)-like cells. Abundance of these cells varied by disease state at 1%, 4%, and 2% of healthy control, non-lesional, and lesional T cells, respectively. Closer inspection revealed that Tfh-like cells from healthy control and non-lesional samples differed in expression of ISGs including *CXCL13* (expressed by 0% of healthy control vs. 76% of non-lesional Tfh-like cells). A similar population of Tfh-like cells was detected in scRNA-seq of kidney biopsies from patients with lupus nephritis^16^, where they were theorized to promote B cell responses such as local antibody production and antigen-specific T cell activation by B cells^17^. Their detection in our dataset suggests a similar process might be occurring in non-lesional skin of lupus patients as well.

Altogether, T cell imbalances and the presence of IFN T cells and other IFN-educated T cell subsets including Tregs in non-lesional samples indicate the presence of an abnormal and likely pathologic T cell infiltrate poised in the prelesional environment of normal-appearing skin of patients with CLE.

### Major shifts in myeloid cell subsets are detected in lesional and non-lesional skin of CLE patients

We then evaluated myeloid cells, the other major immune cell type detected in our skin samples. Sub-clustering and annotation identified nine myeloid cell subsets with largely distinct marker genes (**Fig. 5a-c**): classical type 1 dendritic cell (cDC1; *CLEC9A*, *IRF8*), classical type 2 dendritic cell subset A (cDC2A; *LAMP3* and *CD1B*), classical type 2 dendritic cell subset B (cDC2B; *CLEC10A*, *IL1B*), plasmacytoid dendritic cell (pDCs; *GZMB*, *JCHAIN*), CD16+ dendritic cell (CD16+ DC; *FCGR3A*, *HES1*), Langerhans cell (LC; *CD207*, *CD1A*), lipid-associated macrophage (LAM; *APOE*, *APOC1*), perivascular macrophage (PVM; *CD163*, *SELENOP*), and plasmacytoid dendritic cell-like cell (pDC-like; *PPP1R14A*, *TRPM4*) that was previously described by Villani *et al.*^18^. Myeloid cell subsets showed far greater variability in representation among healthy control, non-lesional, and lesional samples than the above cell types (**Fig. 5a,d,f**).

**Figure 5.**
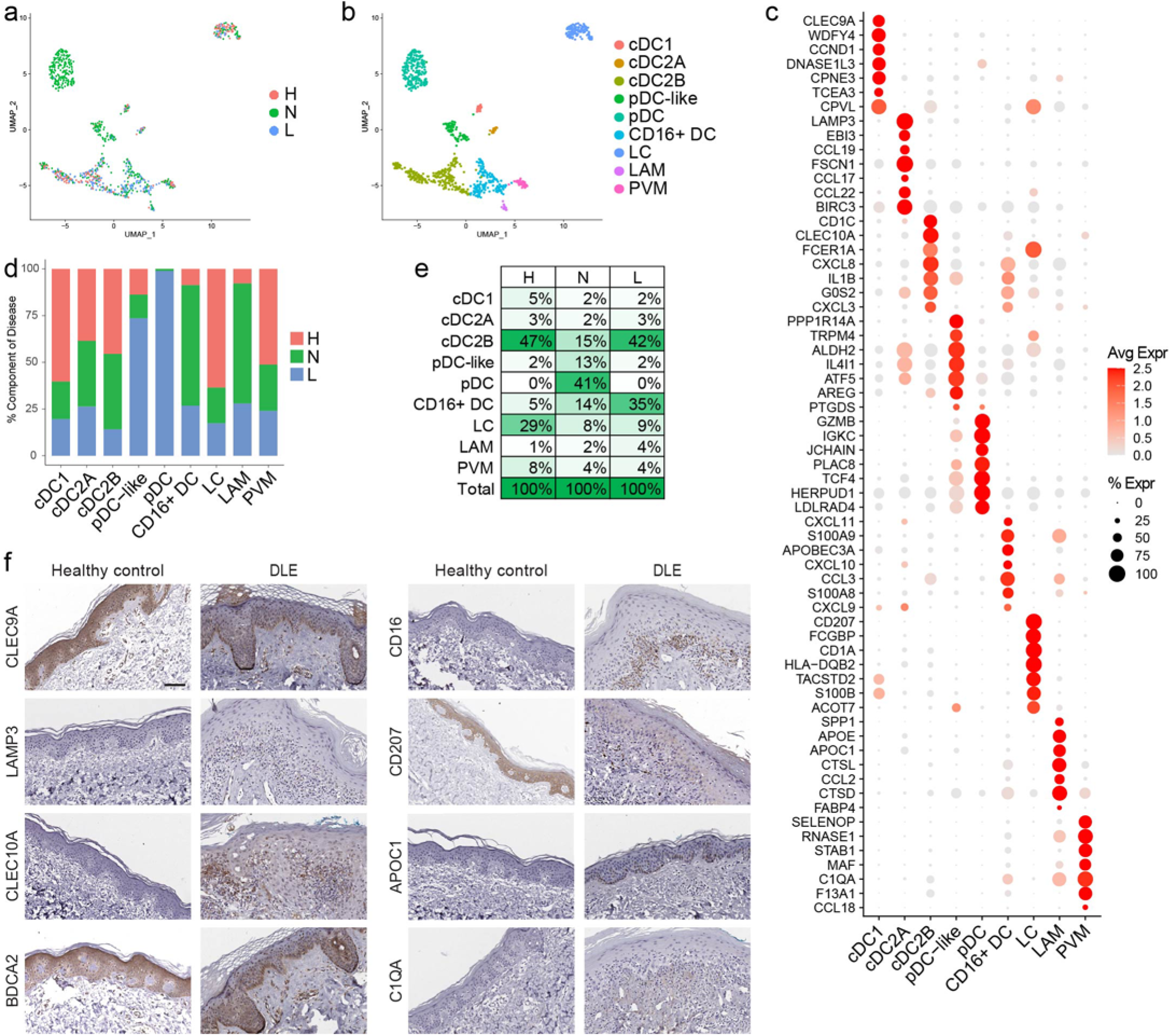
Non-lesional skin of CLE patients shows major myeloid cell subset shifts including infiltration of plasmacytoid and CD16+ dendritic cells. **a.** UMAP plot of 973 myeloid cells colored by disease state. **b.** UMAP plot of myeloid cells colored by subset. cDC1, classical type 1 dendritic cells (DCs); cDC2A, classical type 2 DC subset A; cDC2B, classical type 2 DC subset B; pDC, plasmacytoid DC; DC, dendritic cell; LC, Langerhans cell; LAM, lipid-associated macrophage; PVM, perivascular macrophage. **c.** Dot plot of representative marker genes for each myeloid cell subset. Color scale, average marker gene expression. Dot size, percentage of cells expressing marker gene. **d.** Bar plot of disease state representation within each myeloid cell subset. **e.** Percentage of cells in each myeloid cell subset divided by disease state. **f.** Immunostaining for the indicated marker genes for each myeloid cell subset in healthy control and discoid lupus erythematosus (DLE) lesional skin. *CLEC9A*, cDC1; *LAMP3*, cDC2A; *CLEC10A*, cDC2B; *BDCA2*, pDC; *CD16*, CD16+ DC; *CD207*, LC; *APOC1*, LAM; and *C1QA*, PVM. Scale bar, 50 μm. Images are representative of 2 healthy control and 3 DLE sections.

In healthy control skin, cDC2Bs accounted for nearly half (47%) of the myeloid cells; this differed greatly from non-lesional lupus skin, where pDCs dominated (41%), although most were from a single patient (**Fig. 5e**). In keeping with prior reports, LCs were decreased in lesional^19, 20^ and non-lesional^6^ lupus skin (**Fig. 5d,e**). Among the most striking differences between healthy control and lupus skin, however, was the overrepresentation of CD16+ DCs in both non-lesional and lesional CLE samples compared to healthy skin (**Fig. 5d,e**). While the exact identity of these cells remains somewhat in flux, CD16+ DCs are gaining recognition as a unique DC subset characterized by expression of *FCGR3A*/CD16a that can be detected as a transcriptomically distinct population (labeled DC4 in work by Villani *et al.* profiling circulating mononuclear cell populations in healthy individuals using scRNA-seq^18^). This population is thought to overlap with CD16+ DCs previously described by MacDonald *et al.* as expressing high levels of CD86 and CD40 and possessing potent T cell stimulatory capabilities^21^. CD16+ DCs also exhibit enhanced capacity relative to cDC2Bs for secretion of inflammatory cytokines upon toll-like receptor stimulation^22^, a capacity that is further enhanced in CD16+ DCs isolated from peripheral blood of patients with SLE^23^. Expansion and enhanced function of CD16+ DCs in lupus patients could therefore promote pathogenesis. Based on shared surface marker expression, this subset may also overlap with 6-Sulfo LacNAc-dendritic cells (slanDCs)^24^, a pro-inflammatory myeloid DC subset that has been linked to lupus immunopathogenesis^25, 26^ and is increased in lesional skin of lupus patients^25^.

### Ligand-receptor (L-R) analysis demonstrates lupus-enriched cell-cell interactions prominently involving CD16+ DCs

Following identification of cellular populations, we then sought to understand how cell-cell communication differed in the skin of lupus patients. We thus performed L-R analyses among all major cell populations within healthy control, non-lesional, and lesional skin samples using CellPhoneDB. Each L-R pair was then assigned to the condition in which it showed the highest interaction score, and the number of interactions for each cell type pair was plotted. Few L-R interactions were strongest in healthy control skin, and the majority of these represented KC-KC crosstalk (**Fig. 6a**). Non-lesional skin, in contrast, showed many more interactions (**Fig. 6b**). FBs represented the main ligand-expressing cell type among non-lesional-enriched pairs, but myeloid and endothelial cells (ECs) were also highly interactive. Additionally, eccrine gland cells participated in a high number of interactions in non-lesional skin as expressers of both ligands and receptors, which is of interest given that perieccrine inflammation is a hallmark of CLE. In lesional skin, however, myeloid and ECs were most prominent among cell-cell interactors (**Fig. 6c**).

**Figure 6.**
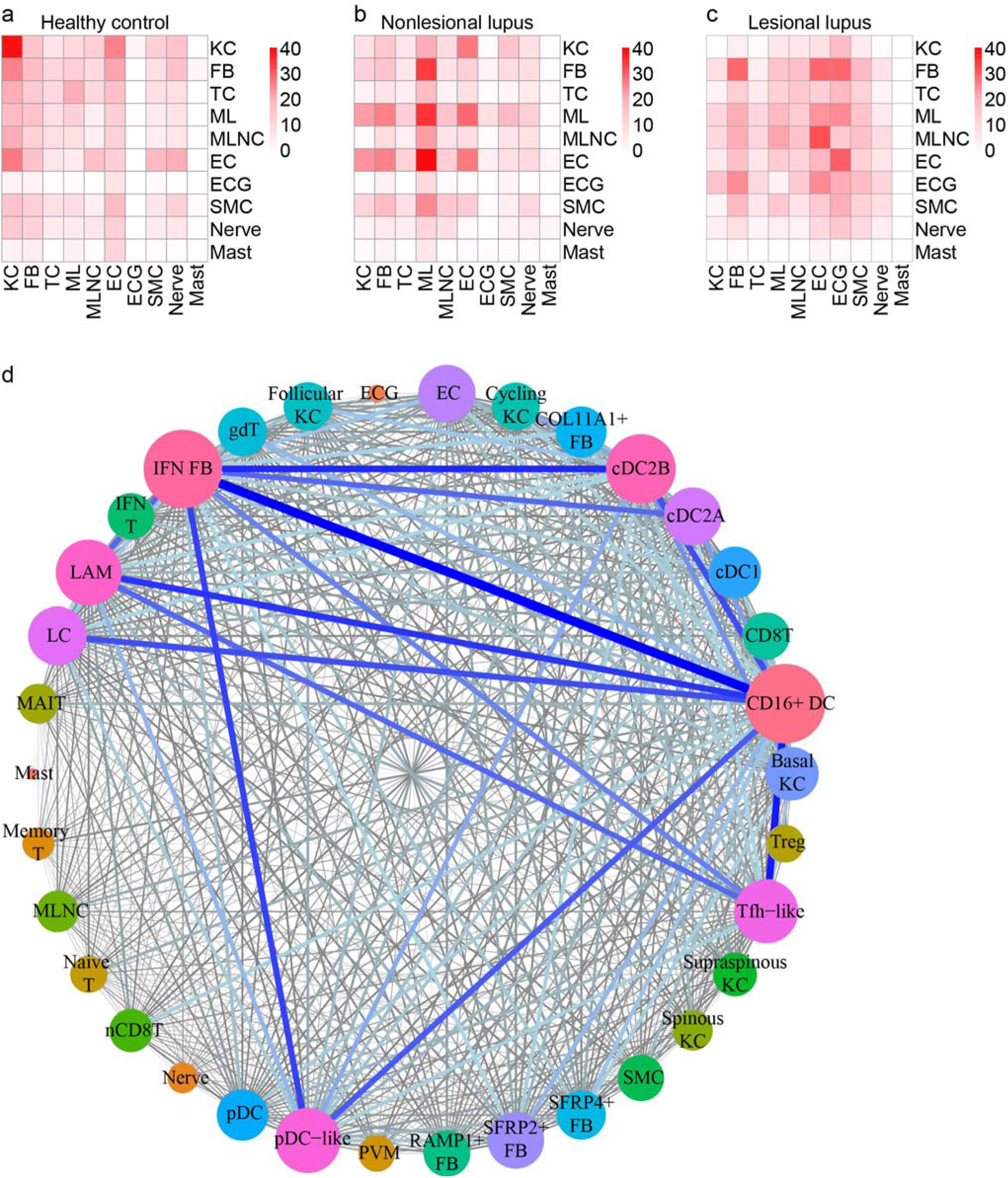
Ligand-receptor (L-R) analysis indicates major shifts in cell-cell communication in CLE and identifies CD16+ DCs as the top candidate cellular interactors in non-lesional skin. **a.** Heatmap depicting the number of L-R pairs with interaction scores highest in H samples divided by cell type. Row, cell type expressing the ligand; column, cell type expressing the receptor. Color scale, number of L-R pairs. **b.** Heatmap depicting the number of L-R pairs with interaction scores highest in N samples. **c.** Heatmap depicting the number of L-R pairs with interaction scores highest in L samples. **d.** Connectome web analysis of interacting cell types. Vertex (colored cell node) size is proportional to the number of interactions to and from that cell type, whereas the thickness of the connecting lines is proportional to the number of interactions between two nodes.

These analyses indicate a prominent role for myeloid cells in non-lesional and lesional skin. Given the functional heterogeneity within the myeloid cell population, we sought to define more precisely the myeloid and other cellular participants in these interactions. Thus, we divided KCs, FBs, T cells, and myeloid cells into their respective subsets and repeated analysis of L-R pairs. Plotting the L-R interactions revealed an even denser network of candidate cellular interactors (**Fig. 6d**). Regarding stromal cells, ligands expressed by KC subsets primarily signaled to receptors on ECs, suggesting a mechanism by which KCs may influence tissue infiltration by immune cells. IFN FBs were among the most active of all cell subsets. However, CD16+ DCs represented the top interactors. Expressing both ligands and receptors, CD16+ DCs showed numerous enriched interactions involving IFN FBs, Tfh-like cells, and cDC2B cells, as well as within-CD16+ DC crosstalk. pDCs were comparatively inert in comparison, rarely participating in L-R pairs (**Fig. 6d**), consistent with recent literature supporting a more senescent phenotype of pDCs in SLE skin^4^.

### Integration of spatial-seq and scRNA-seq analyses provides architectural context shaping cell-cell interactions within lupus skin

L-R analysis derives exclusively from differential gene enrichment and therefore lacks critical spatial context to substantiate putative interactions. To bolster our interaction analyses, we analyzed discoid lupus lesional skin sections using spatial sequencing on the 10x Genomics Visium platform. A section containing multiple hair follicle segments was selected for in depth analysis (**Fig. 7a**). We detected 632 spatially defined spots with an average of 3,704 genes and 10,176 transcripts per spot. Abundant dermal deposition of extracellular glycosaminoglycans, termed mucin, is a frequent feature of cutaneous lupus and is evident in the section as collagen fiber splaying; accordingly, many dermal areas showed very low transcript detection and were excluded during quality control.

**Figure 7.**
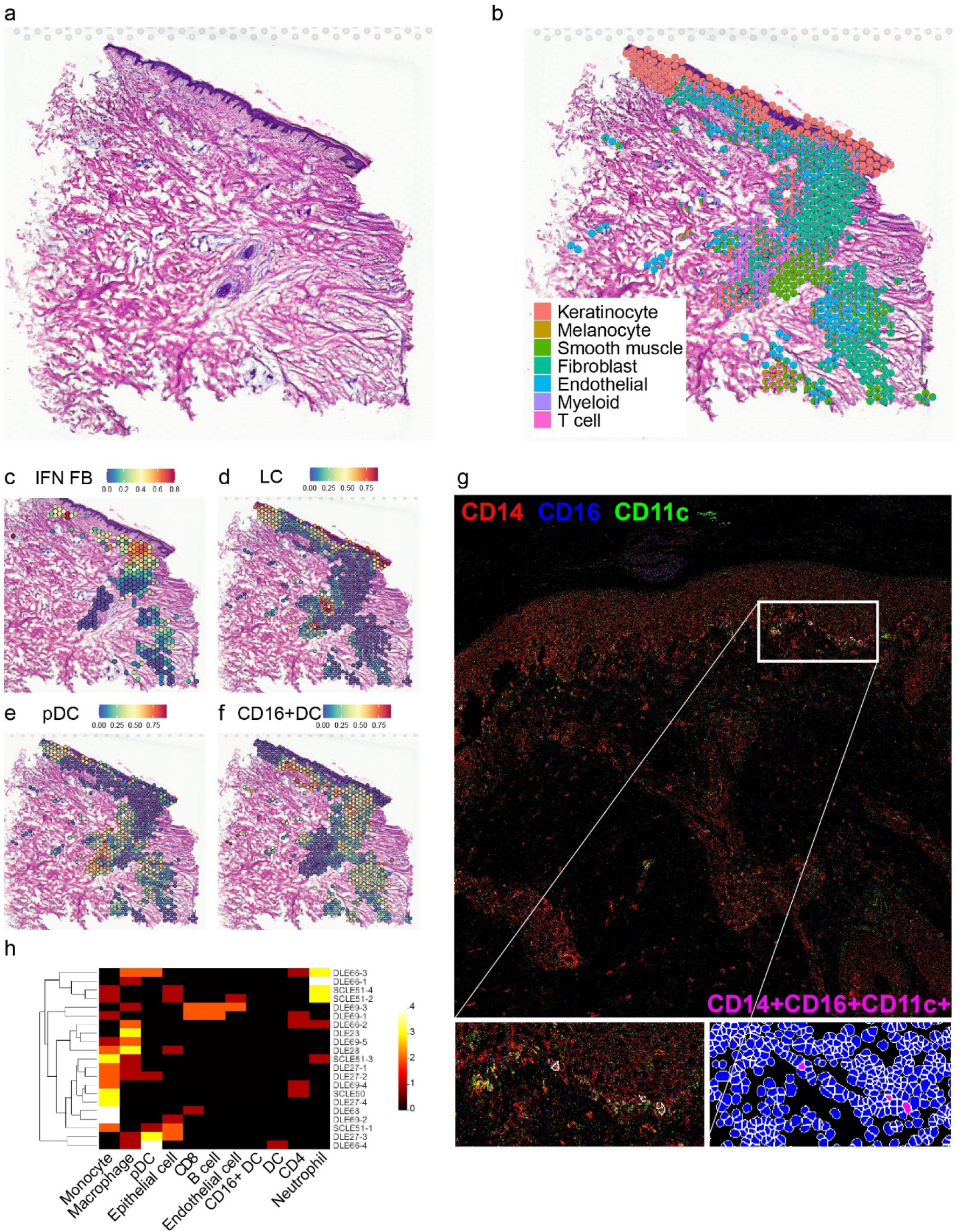
Spatial sequencing reinforces the effects of IFN-producing interfollicular KCs on CD16+ DCs and FBs in the superficial dermis. N=4; data are shown for the most complex sample as defined by the highest number of spots after quality control steps. **a.** Hematoxylin and eosin staining of DLE tissue section corresponding to spatial sequencing data below. **b.** Spatial scatter pie plot showing cell type composition based on detection of scRNA-seq signatures corresponding to 7 cell types. Each spot is represented as a pie chart showing relative cell type proportions. Spot coordinates correspond to tissue location. **c.** Spatial heatmap of the IFN FB subset gene signature. Color, scaled expression of each subset gene signature. Only spots meeting a FB prediction score threshold of 0.25 are shown. **d.** Spatial heatmap of the LC subset gene signature. All spots are shown. **e.** Spatial heatmap of the pDC subset gene signature. All spots are shown. **f.** Spatial heatmap of the CD16+ DC subset gene signature. All spots are shown. **g.** Representative image of a DLE skin section showing the localization of CD16+ DCs (as indicated by the CD14+CD11c+CD16+ immunophenotype) generated by imaging mass cytometry. Insets, subepidermal enrichment of CD16+ DCs. **h.** Heatmap depicting the number of the indicated cell types (columns) located within 4μm of each of the CD16+ DCs (rows) located across 6 DLE (N=16 CD16+ DCs) and 2 subacute CLE (SCLE; N=5CD16+ DCs) sections. Color scale, number of neighboring cells.

At a diameter of 55μm, each spot may account for multiple cells of intermixed types. This was corroborated by spatial heatmaps showing overlapping expression of representative marker genes corresponding to the major cell types (**Supplementary Fig. 3a**). Accordingly, rather than assigning a single cell type to each spot, we generated a pie chart for each spot showing the representation of the transcriptomic signature of each major cell type (**Fig. 7b**, **Supplementary Fig. 3b**). This approach recapitulated the architecture visible on H&E staining, with the epidermis and follicle showing high KC signature detection and the dermis showing a mix of signatures corresponding primarily to FBs, ECs, and smooth muscle cells, which include both vascular smooth muscle and the cells of the arrector pili muscle attached to the hair follicle. The majority of spots with high immune cell signatures localize to the subepidermal and perifollicular regions (**Fig. 7b**), corresponding respectively to the characteristic interface dermatitis and periadnexal infiltrate of discoid lupus. Subepidermal spots showed a particularly prominent myeloid signature with a comparatively weak T cell signature, suggesting strong localization of myeloid cells to the interface, possibly as a direct effect of signaling initiated by KCs or FBs in the prelesional CLE environment.

To further understand how the KC, FB, T cell, and myeloid cell heterogeneity observed in our scRNA-seq mapped onto the architecture of the lesional tissue, we generated spatial heatmaps showing prediction scores corresponding to the subsets defined above. Spatial KC subset analysis demonstrated appropriate localization of the supraspinous, spinous, and basal KC signals to the superficial, mid, and basal epidermis, respectively (**Supplementary Fig. 3c**). The follicular KC signal localized to the follicular epithelium and the cycling KC signal primarily to the deeper portion of the follicle, where the stem cells that give rise to the follicle are located. Consistent with the interfollicular epidermis representing the primary site of exaggerated IFN education in CLE, spatial FB subset analysis revealed prominent localization of the IFN FB signal to the superficial dermis (**Fig. 7c**) and detection of other FB subsets in the perifollicular dermis (**Supplementary Fig. 3d**). Spatial T cell subset analysis also showed subset-specific localization within the tissue section (**Supplementary Fig. 3e**).

Myeloid cell subset spatial heatmaps (**Supplementary Fig. 3f**, full panel) complemented our immunostaining results (**Fig. 5f**). As expected, spots showing a strong LC gene signature were restricted to the epidermis and the follicular epithelium (**Fig. 7d**). Many spots in the perifollicular dermis scored highly for pDCs (**Fig. 7e**). Spots scoring highly for CD16+ DC clustered most densely in the superficial dermis (**Fig. 7f**), again suggesting these cells can be modulated in the IFN-rich environment generated by the basal KCs of the interfollicular epidermis^2^.

To further dissect CD16+ DC cell-cell communication at the level of the individual cell, we performed imaging mass cytometry of lesional DLE and subacute CLE (SCLE) skin biopsies, defining CD16+ DCs as CD14+CD11c+CD16+ cells. This identified CD16+ DCs primarily concentrated in the superficial dermis directly under the dermo-epidermal junction (**Fig. 7g**). Enumeration of neighboring cells revealed that diverse immune cell types are detected within 4 μm distance of CD16+ DCs, with monocytes and macrophages being more common than lymphocytes (**Fig. 7h**). Among stromal cell types included in the analysis, epithelial cells (here, keratinocytes), occurred more commonly in proximity to CD16+ DCs than ECs, supporting interaction between CD16+ DCs and basal KCs. Statistical analysis demonstrated significant overrepresentation of innate inflammatory cells including pDCs and monocytes in proximity to CD16+ DCs in both DLE and SCLE (**Supplementary Fig. 4**).

### Pseudotime analysis of paired circulating and skin-infiltrating myeloid cells suggests that CD16+ DCs arise from non-classical monocytes that undergo IFN education in lupus skin

Overall, our data thus far supported close communication of CD16+ DCs with stromal cells in the skin, so we next wanted to understand the origin and phenotype of CD16+ DCs that infiltrate the skin in lupus patients. Reasoning that these likely arise from circulating mononuclear cells similar to the DC4 subset identified by Villani *et al.*^18^, we examined peripheral blood mononuclear cells (PBMCs) by scRNA-seq from the same seven lupus patients above as well as from four healthy controls. PBMC and skin cell data were aggregated for clustering, and clusters containing myeloid cells were selected for further analysis based on expression of established markers (**Supplementary Fig. 5**). Myeloid cells from PBMCs and skin were then re-clustered together to assess for connections between circulating and skin-infiltrating subsets.

Sub-clustering of the aggregated myeloid cells revealed an apparent transition spanning PBMCs and skin (**Fig. 8a**). Annotation identified the bridging cells as CD14^+^CD16^++^ non-classical monocytes (ncMos), which derived exclusively from PBMCs, and CD16+ DCs from both PBMCs and skin (**Fig. 8b-d**). Pseudotime analysis of ncMos and CD16+ DCs performed using Monocle arranged the cells along a single trajectory reflecting a transition from circulating ncMos to skin-infiltrating CD16+ DC (**Fig. 8e**) – a process that appears to occur much more frequently in lupus than healthy control skin based on relative abundance (**Fig. 8c**). This increased exit into the skin could even account for the decreased proportion of ncMos and circulating CD16+ DCs observed in lupus patients relative to controls in our dataset (**Fig. 8c**).

**Figure 8.**
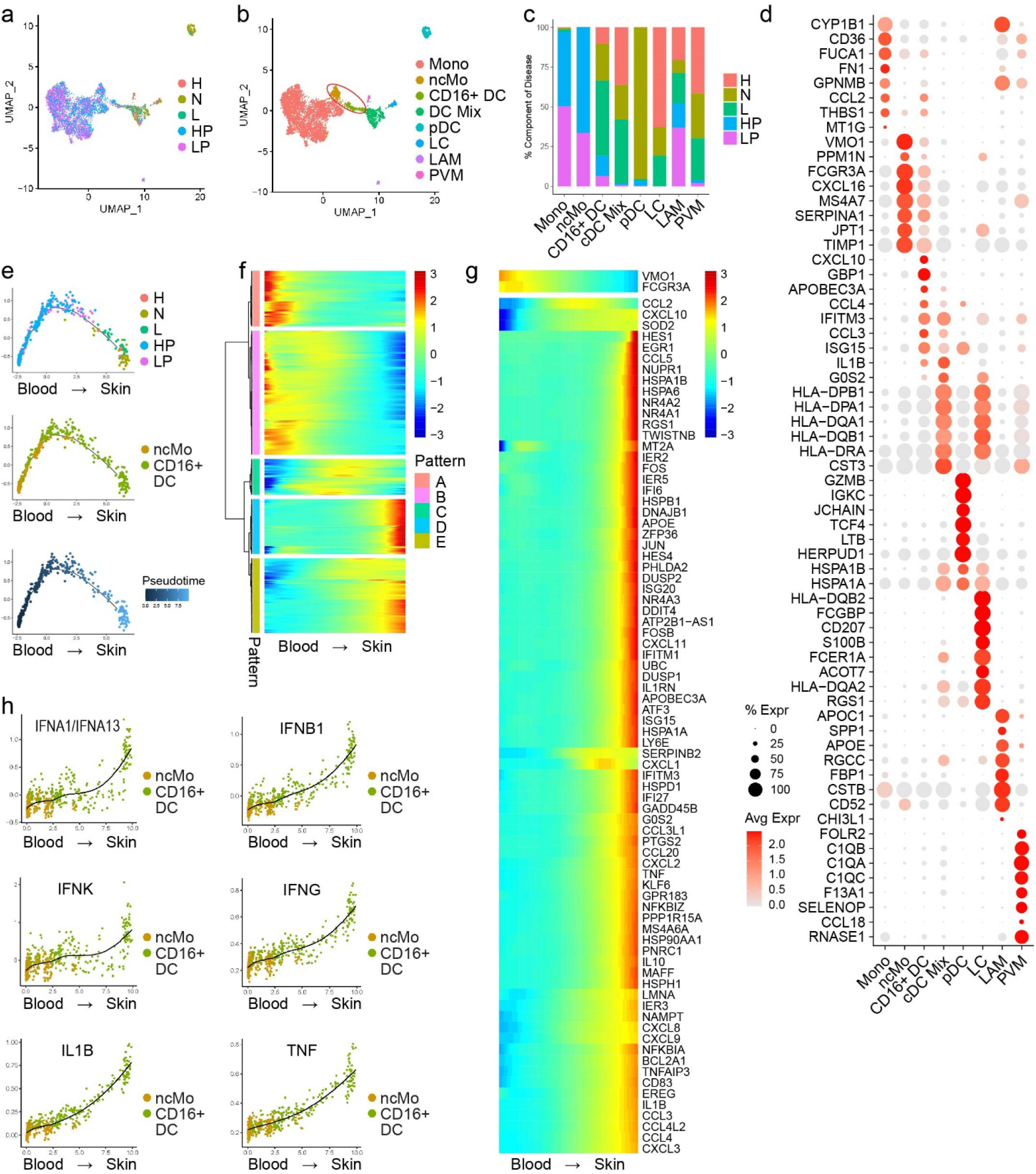
Pseudotime analysis suggests CD16+ DCs arise from non-classical monocytes (ncMos) that migrate into skin of lupus patients and undergo IFN education. **a.** UMAP plot of 6,576 myeloid cells from skin and peripheral blood mononuclear cells (PBMCs) colored by origin and disease state. H, healthy skin; N, non-lesional lupus skin; L, lesional lupus skin; HP, healthy PBMCs; LP, lupus PBMCs. **b.** UMAP plot of myeloid cells colored by subset. Red ellipse, bridge between circulating and skin-derived myeloid cells consisting of non-classical monocyte (ncMo) and CD16+ DC subsets. Mono, monocytes. **c.** Bar plot of origin and disease state representation within each myeloid cell subset. **d.** Dot plot of representative marker genes for each myeloid cell subset. Color scale, average marker gene expression. Dot size, percentage of cells expressing marker gene. **e.** Pseudotime trajectory of ncMos and CD16+ DCs colored by origin and disease state (top), by subset (mid), and by pseudotime (bottom). X-axis, component 1; Y-axis, component 2. **f.** Pseudotime heatmap depicting expression of significant marker genes corresponding to 5 expression patterns that span the transition from ncMo to CD16+ DC. Color scale, scaled marker gene expression across pseudotime. **g.** Pseudotime heatmap depicting expression of the top 80 marker genes across the transition from ncMo to CD16+ DC. Color scale, scaled marker gene expression across pseudotime. **h.** Scatter plots depicting scores for the indicated upstream regulators for each cell across pseudotime. X-axis, pseudotime; Y-axis, module score.

To understand the transcriptional changes that accompany this transition, we performed differential expression analysis along the pseudotime. This identified five gene expression patterns spanning the transition from ncMo to CD16+ DC (**Fig. 8f**). Close inspection of the top 80 most significant marker genes revealed that this transition was marked primarily by late upregulation of numerous genes encoding chemokines, which could support retention of CD16+ DCs and recruitment of other inflammatory cells into the skin of CLE patients, as well as ISGs, consistent with migration of these cells into the type I IFN-rich lupus skin environment (**Fig. 8g**). Broadly, these changes support our prior findings that keratinocyte-derived type I IFN can promote DC activation, enabling them to stimulate immune responses in the skin^2^.

We next analyzed these five gene expression patterns for canonical pathway enrichment for insight into what cellular functions might be acquired and lost across this transition (**Supplementary Fig. 6**). Top canonical pathways enriched among gene patterns expressed earlier in the transition (mostly periphery-associated) tended to relate to leukocyte trafficking. These included ephrin receptor signaling (p=7.44×10^−3^), the top pathway enriched in pattern A, as well as actin cytoskeleton signaling (p=5.57×10^−7^) and integrin signaling (p=4.57×10^−4^), the top pathways enriched in pattern B. Top pathways enriched among gene patterns expressed later in the transition (mostly skin-associated) related more to cytokine signaling. The top pathway enriched in pattern D was the role of hypercytokinemia/hyperchemokinemia in the pathogenesis of influenza (p=6.79×10^−6^), with IFN signaling (p=2.80×10^−4^) also highly ranked. The top pathway enriched in pattern E was IL-6 (p=1.85×10^−6^), in line with data supporting a role for increased production of IL-6 by lupus KCs secondary to conditioning by IFN signaling^3^.

To analyze the cytokines influencing the transition from ncMo to skin-infiltrating CD16+ DC, we calculated upstream regulator scores for all cytokines included in IPA for each cell and determined the correlation of these scores with pseudotime. Scores for a panel of type I IFNs (IFN-α1/13, IFN-β, and IFN-Κ) and IFN-γ (**Fig. 8h**) indicated very high correlation (r= 0.774, 0.880, 0.666, and 0.873, respectively) with a more pronounced rise late in pseudotime, consistent with robust IFN education representing an important terminal step in this transition. A number of upstream regulators showed more gradual induction across pseudotime and higher correlation scores, suggestive of an earlier role in the transition. These included IL-1β (r= 0.928), the top-correlated cytokine, and TNF (r= 0.903); of note, these also emerged as top significant genes in the pseudotime DEG analysis, showing late upregulation in the transition from ncMo to CD16+ DC (**Fig. 8g**). This is in keeping with our prior report that prolonged type I IFN exposure primes monocytes for inflammasome activation and enhances their production of IL-1β^27^.

The interactions that mediate infiltration of CD16+ DCs into normal-appearing skin of patients with CLE are not yet known. To highlight L-R pairs that could promote this accumulation, we generated circos plots of all cytokine interaction pairs in which CD16+ DCs expressed either receptor (**Supplementary Fig. 7a**) or ligand (**Supplementary Fig. 7b**). CD16+ DCs expressed 12 cytokine receptors involved in L-R pairs (**Supplementary Fig. 7a**). ECs and smooth muscle cells, which include vascular smooth muscle cells, expressed the highest number of interacting ligands. Next highest among stromal cells were IFN FBs. In a similar pattern, CD16+ DCs expressed 23 ligands involved in L-R pairs (**Supplementary Fig. 7b**), with ECs and IFN FBs expressing the highest number of interacting receptors. Together, these L-R data suggest enhanced interactions with ECs and IFN FBs enable CD16+ DCs to accumulate in the skin of CLE patients, where the IFN-rich environment augments their pro-inflammatory properties and capacity for cell-cell communication.

## DISCUSSION

Collectively, our data describe the cellular composition and architecture of cutaneous lupus at unprecedented resolution. We demonstrate the pervasive effects of IFN, of which the epidermis is a critical source^2^, on skin stromal and immune cells alike. Most intriguingly, these effects are pronounced in non-lesional samples, suggesting that normal-appearing skin of patients with CLE exists in an immunologically primed, ‘prelesional’ state. This state skews the transcriptional programs of many of the major cell types in the skin, with dramatic effects on the capacity for cell-cell communication. Indeed, even the minor cellular constituents of the skin not examined in detail here exhibited transcriptional shifts in non-lesional CLE skin that alter their potential to engage other stromal and immune cells (**Fig. 6b**).

This investigation highlighted CD16+ DCs, a myeloid cell subset increasingly implicated in lupus pathogenesis^21, 22, 23, 25, 26^, as proficient intercellular communicators even in non-lesional skin of CLE patients (**Fig. 6d**), where they are highly abundant (**Fig. 5d**). Unsupervised clustering followed by pseudotime analysis of combined myeloid cells from skin and peripheral blood suggests that progenitor ncMos in circulation give rise CD16+ DCs (**Fig. 8a,b,e**). Clustering of DCs isolated from peripheral blood of healthy patients identified a subset of so-called DC4 cells characterized by expression of *FCGR3A*, encoding CD16a, with high transcriptional similarity to a monocyte subset with features of ncMos^18^, inspiring some discussion that DC4 cells may in fact represent monocytes^28, 29^. Here, however, utilization of single-cell technologies across blood and tissues has enabled identification of a CD16+ DC population enriched in the skin of patients with SLE and defined their probable precursors and the transcriptomic changes accompanying this transition that arm CD16+ DCs to instigate tissue inflammation. ScRNA-seq analysis of immune cells isolated from kidney biopsies of patients with lupus nephritis (LN) and healthy controls suggests that this paradigm may not be limited to the skin. Arazi *et al.* identified several myeloid subpopulations resembling DC4 cells (CM0, CM1, and CM4); these were highly enriched in LN biopsies and showed upregulation of IFN response scores relative to steady-state kidney macrophages and conventional DCs^16^, suggesting IFN education is a characteristic feature of tissue-infiltrating CD16+ DCs that facilitates their pathogenicity in lupus.

Accumulation of CD16+ DCs in the skin represents a distinguishing feature of not just the lesional but also the prelesional CLE environment. In CLE patients, exit of CD16+ DCs from the circulation into non-lesional skin is likely enhanced through robust L-R interactions between CD16+ DCs and the vasculature (**Supplementary Fig. 7**). Following tissue infiltration, CD16+ DCs may be directed to accumulate in the superficial dermis (**Fig. 7g**) by ligand gradients generated by subepidermal FBs that have been transformed to an IFN FB phenotype through the effects of KC-secreted IFN (**Fig. 3g**, **Fig. 7c**). Upon encountering the IFN-rich non-lesional CLE environment, CD16+ DCs upregulate a panoply of ISG-encoded cytokines and chemokines (**Fig. 8f,g**). This endows them with the capacity for extensive cell-cell communication with diverse cell types (**Supplementary Fig. 7**) including innate immune cells in their immediate proximity (**Supplementary Fig. 4**), whereby they may contribute to genesis of CLE lesions. Thus, the prelesional environment of skin of CLE patients represents a collaboration between stromal and immune cells, with critical contributions from KCs, FBs, and CD16+ DCs. Further investigations directed at differences in cell-cell communication between non-lesional and lesional CLE will provide additional insight into the downstream events that precipitate clinically evident inflammation and lesion formation and provide targeted therapeutic strategies to improve treatment response in patients with this devastating disease.

## METHODS

### Human sample acquisition

7 patients with active CLE (**Supplementary Table 1**) were recruited for this study, all of whom contributed lesional and non-lesional (sun-protected skin of the buttock) 6mm punch skin biopsies and whole blood for isolation of PBMCs. A diagnosis of SLE was confirmed for 6 of 7 patients via the European League Against Rheumatism/American College of Rheumatology criteria^30^. 14 healthy controls were recruited for skin biopsy and 4 for whole blood. The study was approved by the University of Michigan Institutional Review Board (IRB), and all patients were consented. The study was conducted according to the Declaration of Helsinki Principles.

### Immunohistochemistry

Paraffin embedded tissue sections from punch biopsies from patients with discoid lupus erythematosus and healthy control skin were heated at 60°C for 30 minutes, de-paraffinized, and rehydrated. Slides were placed in antigen retrieval buffer at the pH indicated in **Supplementary Table 3** and heated at 125°C for 30 seconds in a pressure cooker water bath. After cooling, slides were treated with 3% H2O2 (5 minutes) and blocked using 10% goat serum (30 minutes). Overnight incubation was performed at 4°C with primary antibodies at the indicated concentrations (**Supplementary Table 3**). Slides were then washed, treated with appropriate secondary antibodies, peroxidase (30 minutes), and diaminobenzidine substrate, before imaging.

### Single-cell RNA library preparation, sequencing, and alignment

Generation of single-cell suspensions for scRNA-seq was performed as follows: Skin biopsies were incubated overnight in 0.4% dispase (Life Technologies) in Hank’s Balanced Saline Solution (Gibco) at 4°C. Epidermis and dermis were separated. Epidermis was digested in 0.25% Trypsin-EDTA (Gibco) with 10U/mL DNase I (Thermo Scientific) for 1 hour at 37°C, quenched with FBS (Atlanta Biologicals), and strained through a 70μM mesh. Dermis was minced, digested in 0.2% Collagenase II (Life Technologies) and 0.2% Collagenase V (Sigma) in plain medium for 1.5 hours at 37°C, and strained through a 70μM mesh. Epidermal and dermal cells were combined in 1:1 ratio, and libraries were constructed by the University of Michigan Advanced Genomics Core on the 10X Chromium system with chemistry v3. Libraries were then sequenced on the Illumina NovaSeq 6000 sequencer to generate 150 bp paired end reads. Data processing including quality control, read alignment (hg38), and gene quantification was conducted using the 10X Cell Ranger software. The samples were then merged into a single expression matrix using the cellranger aggr pipeline.

### Cell clustering and cell type annotation

The R package Seurat (v3.1.2)^31^ was used to cluster the cells in the merged matrix. Cells with less than 500 transcripts or 100 genes or more than 10% of mitochondrial expression were first filtered out as low-quality cells. The NormalizeData function was used to normalize the expression level for each cell with default parameters. The FindVariableFeatures function was used to select variable genes with default parameters. The ScaleData function was used to scale and center the counts in the dataset. Principal component analysis (PCA) was performed on the variable genes, and the first 30 PCs were used in the RunHarmony function from the Harmony package^32^ to remove potential batch effect among samples processed in different libraries. Uniform Manifold Approximation and Projection (UMAP) dimensional reduction was performed using the RunUMAP function. The clusters were obtained using the FindNeighbors and FindClusters functions with the resolution set to 0.5. The cluster marker genes were found using the FindAllMarkers function. The cell types were annotated by overlapping the cluster markers with the canonical cell type signature genes. To calculate the disease composition based on cell type, the number of cells for each cell type from each disease condition were counted. The counts were then divided by the total number of cells for each disease condition and scaled to 100 percent for each cell type. Differential expression analysis between H and N or H and L were carried out using the FindMarkers function.

### Cell type sub-clustering

Sub-clustering was performed on the abundant cell types. The same functions described above were used to obtain the sub-clusters. Sub-clusters that were defined exclusively by mitochondrial gene expression, indicating low quality, were removed from further analysis. The subtypes were annotated by overlapping the marker genes for the sub-clusters with the canonical subtype signature genes. Ingenuity pathway analysis was applied to the differentially expressed genes to determine the canonical pathways and the potential upstream regulators. The upstream regulators with an activation z score ≥2 or ≤2 were considered significant. The module scores were calculated using the AddModuleScore function on the genes activated by the intended cytokine from bulk RNA-seq analysis as previously described^33^.

### Ligand receptor interaction analysis

CellphoneDB (v2.0.0)^34^ was applied for L-R analysis. In the first analysis, the major cell type annotations were used. The cells were separated by their disease classifications (H, N, L), and a separate run was performed for each disease classification. Pairs with p value >0.05 were filtered out from further analysis. To compare among the three disease conditions, each pair was assigned to the condition in which it showed the highest interaction score. The number of interactions for each cell type pair was then calculated. In the second analysis, KCs, FBs, T cells, and myeloid cells were divided into their respective subtypes. The number of interactions between each cell type pair was calculated. The cytokine pairs for CD16+ DC was plotted in circos plots using the R package “circlize”. The connectome web was plotted using the R package “igraph”.

### Pseudotime trajectory construction

Pseudotime trajectory for myeloid cell sub-clusters 6 and 8 from **Fig. 8a** was constructed using the R package Monocle (v2.10.1)^35^. The raw counts for cells were extracted from the Seurat analysis and normalized by the estimateSizeFactors and estimateDispersions functions with the default parameters. Genes detected in >10 cells were retained for further analysis. Variable genes were determined by the differentialGeneTest function with a model against the Seurat sub-cluster identities. The orders of the cells were determined by the orderCells function, and the trajectory was constructed by the reduceDimension function with default parameters. Differential expression analysis was carried out using the differentialGeneTest function with a model against the pseudotime, and genes with an adjusted p value smaller than 0.05 were clustered into five patterns and plotted in the heatmap. Ingenuity Pathway Analysis was used to determine the upstream regulators for the genes in each expression pattern. A module score was calculated for each upstream regulator on gene targets from all five patterns. The module scores were calculated using the Seurat function AddModuleScore with default parameters. Pearson correlation was then performed between the upstream regulator module scores and the pseudotime.

### Spatial sequencing library preparation

Skin samples were frozen in OCT medium and stored at −80°C until sectioning. Optimization of tissue permeabilization was performed on 20 μm sections using Visium Spatial Tissue Optimization Reagents Kit (10X Genomics, Pleasanton, CA, USA), which established an optimal permeabilization time to be 9 minutes. Samples were mounted onto a Gene Expression slide (10X Genomics), fixed in ice-cold methanol, stained with hematoxylin and eosin, and scanned under a microscope (Keyence, Itasca, IL, USA). Tissue permeabilization was performed to release the poly-A mRNA for capture by the poly(dT) primers that are precoated on the slide and include an Illumina TruSeq Read, spatial barcode, and unique molecular identifier (UMI). Visium Spatial Gene Expression Reagent Kit (10X Genomics) was used for reverse transcription to produce spatially barcoded full-length cDNA and for second strand synthesis followed by denaturation to allow a transfer of the cDNA from the slide into a tube for amplification and library construction. Visium Spatial Single Cell 3□ Gene Expression libraries consisting of Illumina paired-end sequences flanked with P5/P7 were constructed after enzymatic fragmentation, size selection, end repair, A-tailing, adaptor ligation, and PCR. Dual Index Kit TT Set A (10X Genomics) was used to add unique i7 and i5 sample indexes and generate TruSeq Read 1 for sequencing the spatial barcode and UMI and TruSeq Read 2 for sequencing the cDNA insert, respectively.

### Spatial sequencing data analysis

After sequencing, the reads were aligned to the human genome (hg38), and the expression matrix was extracted using the spaceranger pipeline. Seurat was then used to analyze the expression matrix. Specifically, the SCTransform function was used to scale the data and find variable genes with default parameters. PCA and UMAP were applied for dimensional reduction. The FindTransferAnchors function was used to find a set of anchors between the spatial-seq data and scRNA-seq data, which were then transferred from the scRNA-seq to the spatial-seq data using the TransferData function. These two functions construct a weights matrix that defines the association between each query cell and each anchor. These weights sum to 1 and were used as the percentage of the cell type in the spots.

### Data availability

The scRNA-seq data are available in GEO under accession number [to be deposited].

### Imaging mass cytometry of tissue sections

Formalin-fixed, paraffin-embedded (FFPE) skin biopsy tissue sections from lesional skin of patients with SCLE or DLE were analyzed using the Hyperion imaging CyTOF system (Fluidigm) as previously described^33^ with modifications of the antibody panel. Specifically, metal-tagged antibodies including pan-keratin (C11, Biolegend), BDCA2 (Polyclonal, R&D Systems), CD56 (123C3, ThermoFisher Scientific), HLA-DR (LN3, Biolegend), CD11c (EP1347Y, Abcam), and CD4 (EPR6855, Fluidigm) were added in this study. Markers used to annotate each cell type are listed in **Supplementary Table 4**.

### Imaging mass cytometry data analysis

Multiplexed imaging mass cytometry data were converted first to .TIFF images using MCD Viewer v1.0.560.2 (Fluidigm) and then segmented using CellProfiler v3.1.8 for single-cell analysis. The unsupervised clustering algorithm Phenograph was performed on 16 markers (CD56, CD15, CD11c, CD16, CD14, CD31, CD27, CD4, HLA-DR, CD68, CD20, CD8, BDCA2, CD138, E-cadherin, and Pan-Keratin) using histoCAT v1.75 software. To identify the rare CD16+ DCs, manual gating was performed on high CD14+CD11c+CD16+ expression. Neighborhood analysis was performed by permutation test^36^ using histoCAT.

## AUTHOR CONTRIBUTIONS

Conceived and designed the analysis: ACB, FM, JEG, JMK.

Collected the data: ACB, OP, MGK, GH, XX, CMY, SMR.

Performed the analysis: FM, RW, CMY, FW, LCT, MP, RLM.

Wrote the paper: ACB, FM, JEG, JMK.

Supervised the work: FW, LCT, MP, RLM, JEG, JMK.

## ACKNOWLEDGMENTS

This work was supported by the Taubman Institute via Innovative Program (JMK, JEG, and FW) and Parfet Emerging Scholar (JMK) funds; the Babcock Endowment Fund (LCT, JEG); and the National Institute of Health (NIH): K08-AR078251 (ACB), R01-AR071384 (JMK), K24-AR076975 (JMK), R01-AR069071 (JEG), R01-AR073196 (JEG), P30-AR075043 (JEG), K01-AR072129 (LCT), R01-AI022553 (RLM), R01-AR040312 (RLM), R01-AR074302 (RLM), R01-AR074302 (RLM), Office of the Director S10-OD020053 (FW), the National Cancer Institute P30-CA046592 (FW), and the Lupus Research Alliance (JMK). ACB is supported by the Dermatology Foundation. LCT is supported by the Dermatology Foundation, Arthritis National Research Foundation, and National Psoriasis Foundation. FW is supported by the National Science Foundation (1653611).

## DECLARATION OF INTERESTS

JMK has received Grant support from Q32 Bio, Celgene/BMS, Ventus Therapeutics, and Janssen. JEG has received Grant support from Celgene/BMS, Janssen, Eli Lilly, and Almirall. JMK has served on advisory boards for AstraZeneca, Eli Lilly, GlaxoSmithKline, Bristol Myers Squibb, Avion Pharmaceuticals, Provention Bio, Aurinia Pharmaceuticals, Ventus Therapeutics, and Boehringer Ingelheim. JEG has served on advisory boards for AstraZeneca, Sanofi, Eli Lilly, Boehringer Ingelheim, Novartis, Janssen, Almirall, BMS. All other authors have nothing to disclose.

## SUPPLEMENTARY FIGURES

**Supplementary Figure 1.**
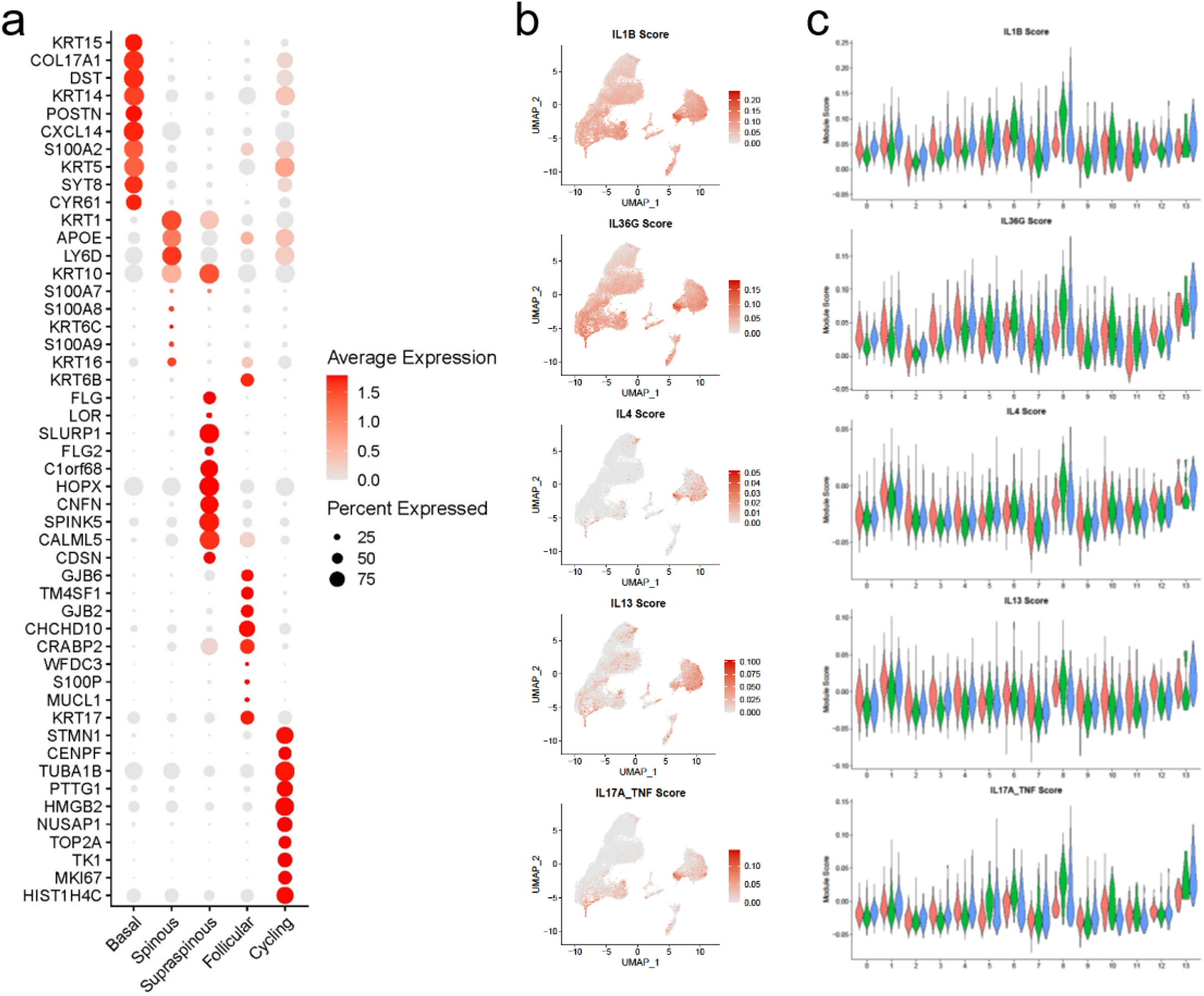
Cytokine module scoring reveals that lupus keratinocyte (KC) transcriptomes are heavily influenced by interferon (IFN). **a.** Dot plot of marker genes showing the highest fold change for each KC subtype. Color scale, average marker gene expression. Dot size, percentage of cells expressing marker gene. **b.** Feature plots of KC scores for the indicated additional cytokine modules. **c.** Violin plots of KC scores for the indicated additional cytokine modules split by sub-cluster and disease state.

**Supplementary Figure 2.**
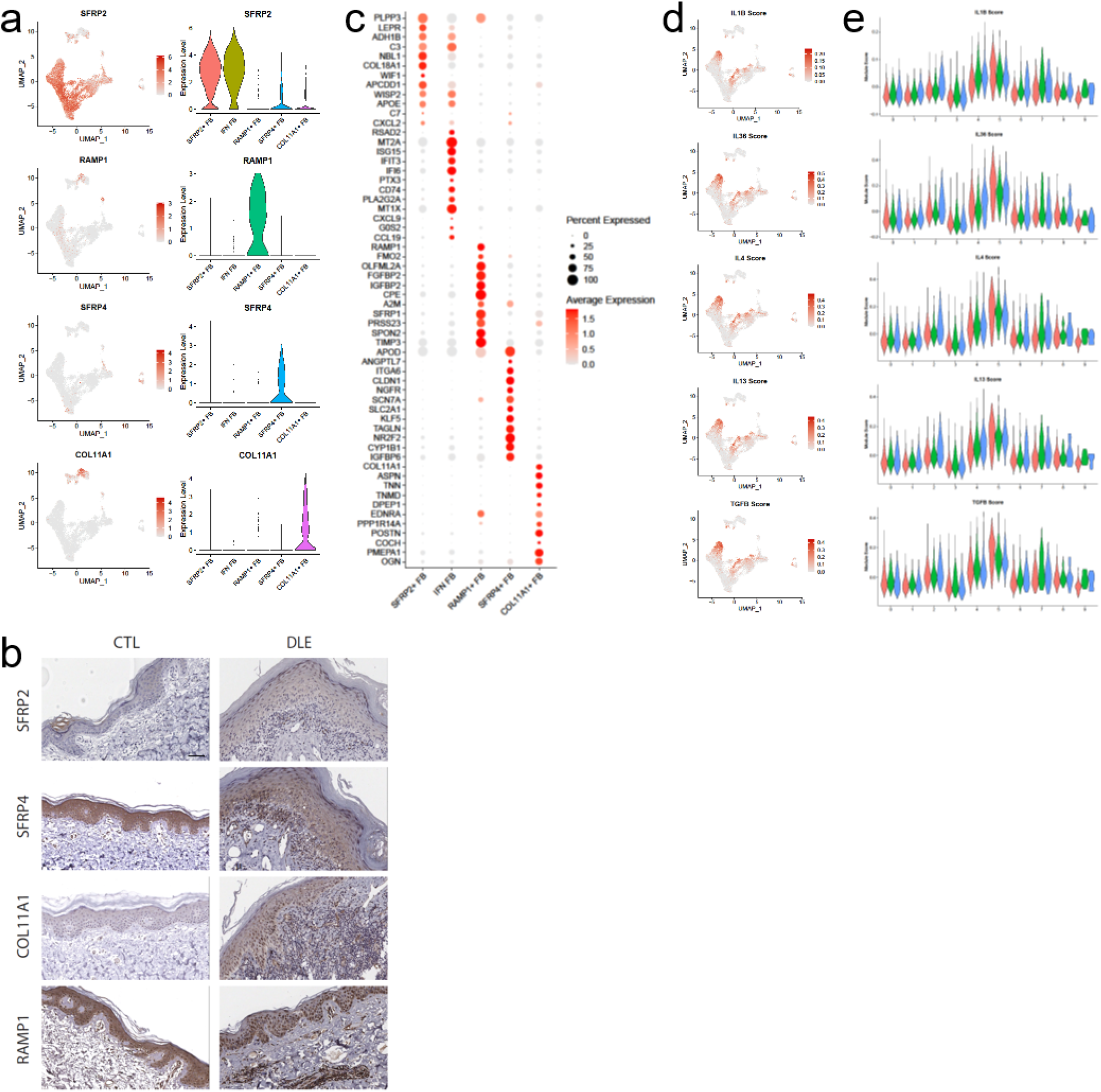
Cytokine module scoring reveals that lupus fibroblast (FB) transcriptomes are heavily influenced by IFN. **a.** Feature and violin plots of the indicated FB subtype markers. **b.** Detection of the indicated FB subtype markers by immunohistochemistry (IHC) in healthy control (CTL) and lesional discoid lupus erythematosus (DLE) skin sections. Scale bar, 50 μm. Images are representative of sections from 2 control CTL and 3 DLE biopsies examined. **c.** Dot plot of marker genes showing the highest fold change for each FB subtype. Color scale, average marker gene expression. Dot size, percentage of cells expressing marker gene. **d.** Feature plots of KC scores for the indicated additional cytokine modules. **e.** Violin plots of KC scores for the indicated additional cytokine modules split by sub-cluster and disease state.

**Supplementary Figure 3.**
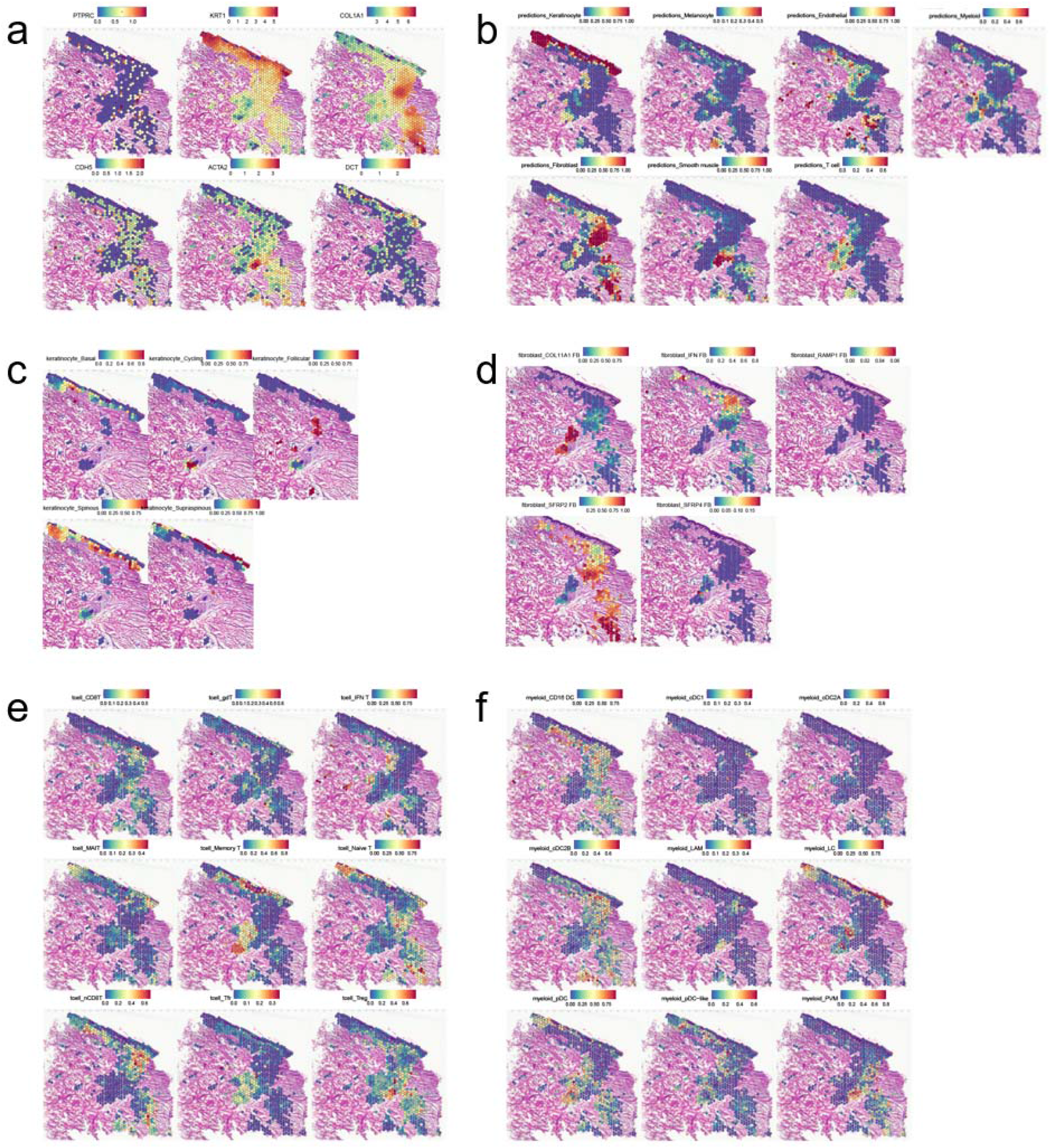
Spatial sequencing enables identification of individual cell types and cell subsets within lesional DLE sections. N=4; data are shown for the most complex sample as defined by the highest number of spots after quality control steps. Full panels are shown, including panels presented in **Fig. 7**. **a.** Spatial heatmap of marker genes identifying immune cells (*PTPRC*, encoding CD45), KCs (*KRT1*), FBs (*COL1A1*), EC (*CDH5*), smooth muscle cells (*ACTA2*), and melanocytes (*DCT*). Color, scaled expression of each gene. Spot coordinates correspond to tissue location. **b.** Spatial heatmaps showing detection of scRNA-seq cell type-specific gene signatures for seven cell types. Color, scaled expression of each cell type signature. **c.** Spatial heatmap of KC subset gene signatures. Only spots meeting a KC prediction score threshold of 0.25 are shown. Color, scaled expression of each subset signature. **d.** Spatial heatmap of FB subset gene signatures. Only spots meeting a FB prediction score threshold of 0.25 are shown. **e.** Spatial heatmap of T cell subset gene signatures. All spots are shown. **f.** Spatial heatmap of myeloid cell subset gene signatures. All spots are shown.

**Supplementary Figure 4.**
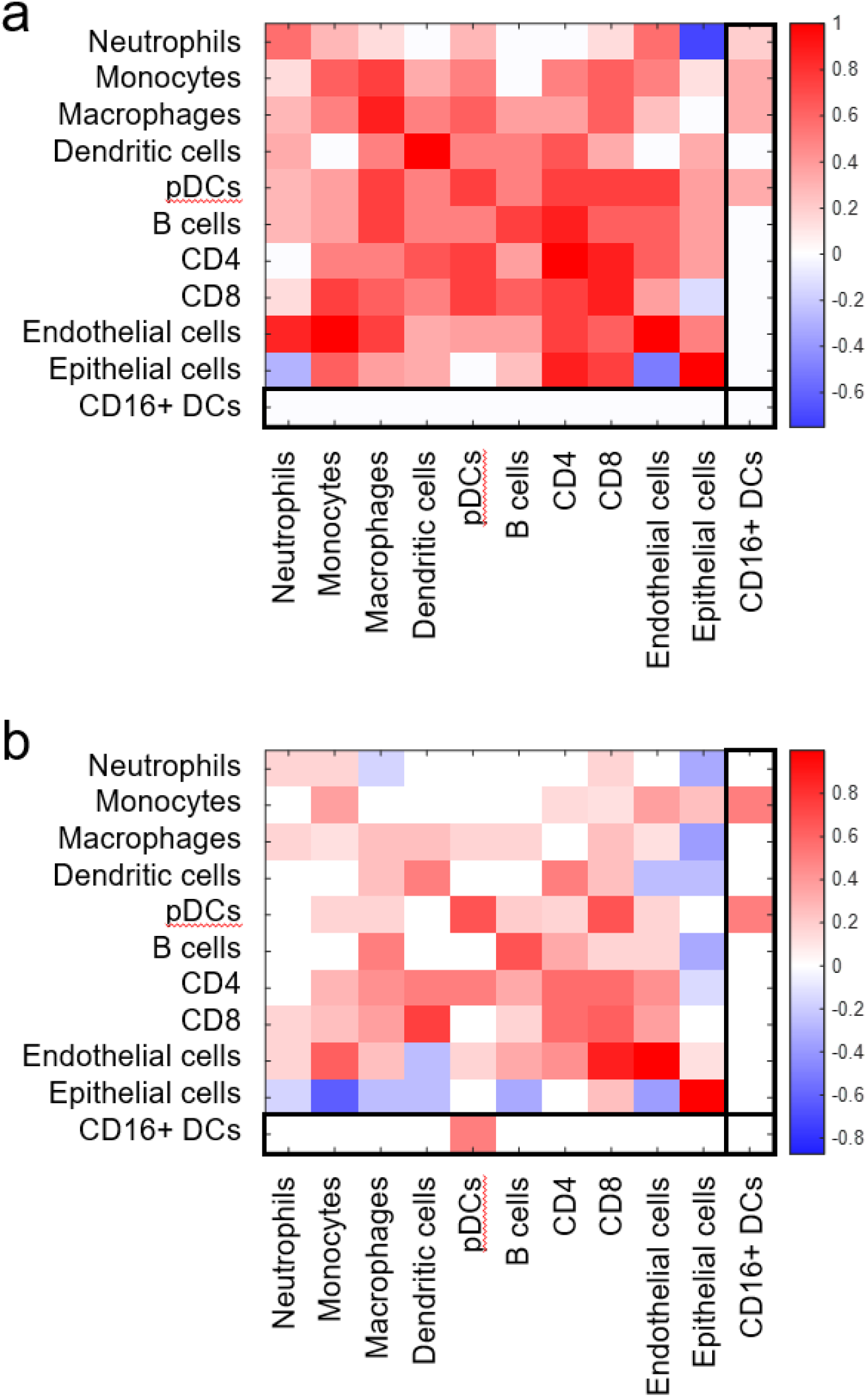
Neighborhood analysis of imaging mass cytometry data reveals CD16+ dendritic cells (DCs) tend to neighbor pDCs and monocytes. Only values for cell pairs with p < 0.05 by permutation test are shown. Values for CD16+ DCs (as indicated by the CD14+CD11c+CD16+ immunophenotype) are boxed. Row, cell type of interest; column, cell type in neighborhood. Color scale, percent of images showing interaction (red) or avoidance (blue). **a.** Neighborhood analysis of the 6 DLE sections analyzed in **Fig. 7h**. **b.** Neighborhood analysis of the 2 SCLE sections analyzed in **Fig. 7h**.

**Supplementary Figure 5.**
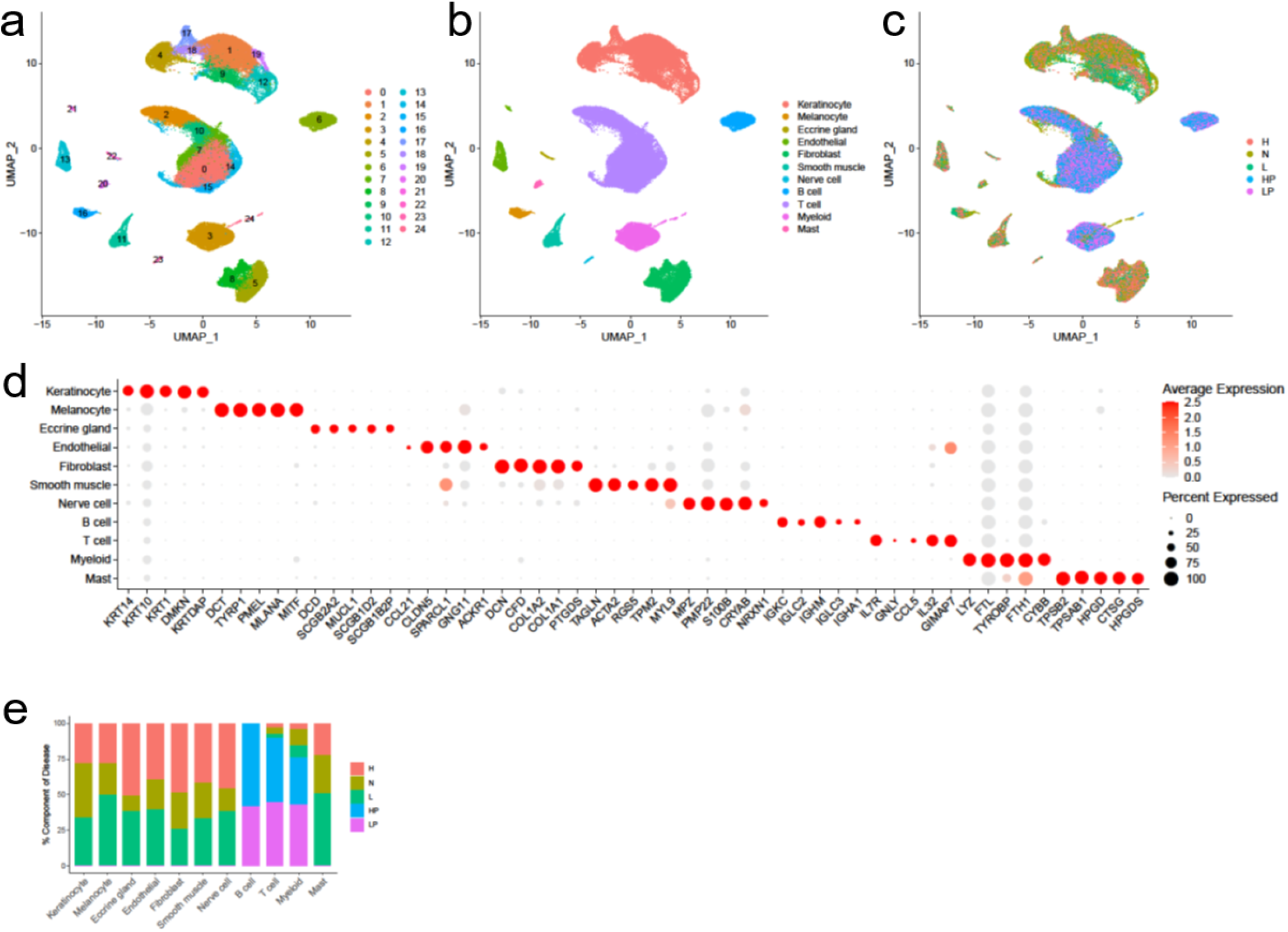
Integrated scRNA-seq of peripheral blood mononuclear cells and skin cells of CLE patients. **a.** UMAP plot of 100,822 cells colored by sub-cluster. **b.** UMAP plot of cells colored by cell type. **c.** UMAP plot of cells colored by origin and disease state. HP, healthy peripheral blood mononuclear cells (PBMCs); LP, lupus PBMCs. **d.** Dot plot of representative marker genes for each cell type. Color scale, average marker gene expression. Dot size, percentage of cells expressing marker gene. **e.** Bar plot of cell type abundance across disease states.

**Supplementary Figure 6.**
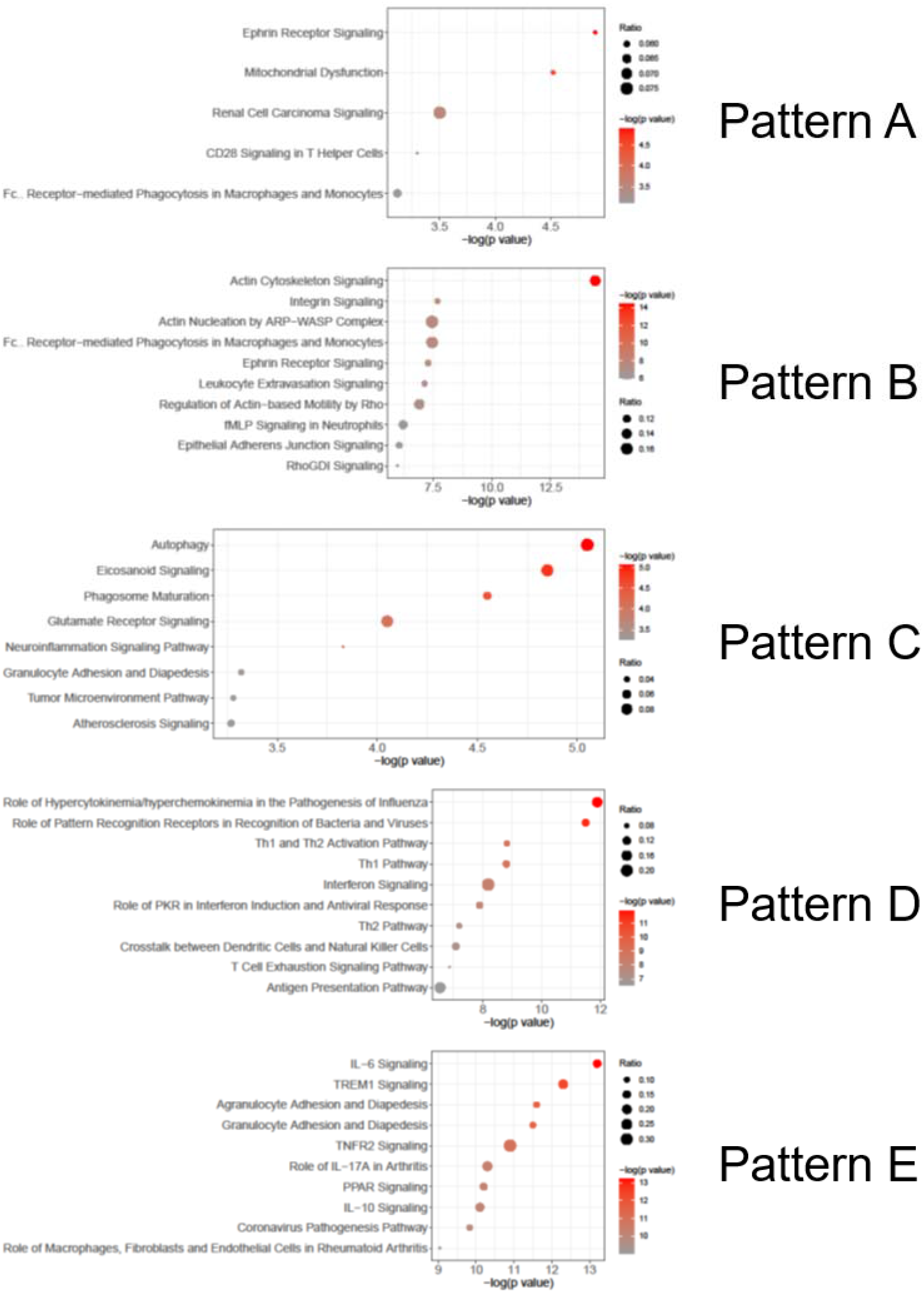
Dot plots showing the top canonical pathways enriched among the gene markers of the five expression patterns across pseudotime. Patterns correspond to the expression heatmap shown in **Fig. 8f**. The top ten pathways (or the total number of significant pathways if fewer than ten) are shown for each pattern. Color scale, log(p value) from the enrichment analysis. Dot size, ratio calculated by dividing the number of associated genes found in the gene marker list by the total number of genes in each pathway.

**Supplementary Figure 7.**
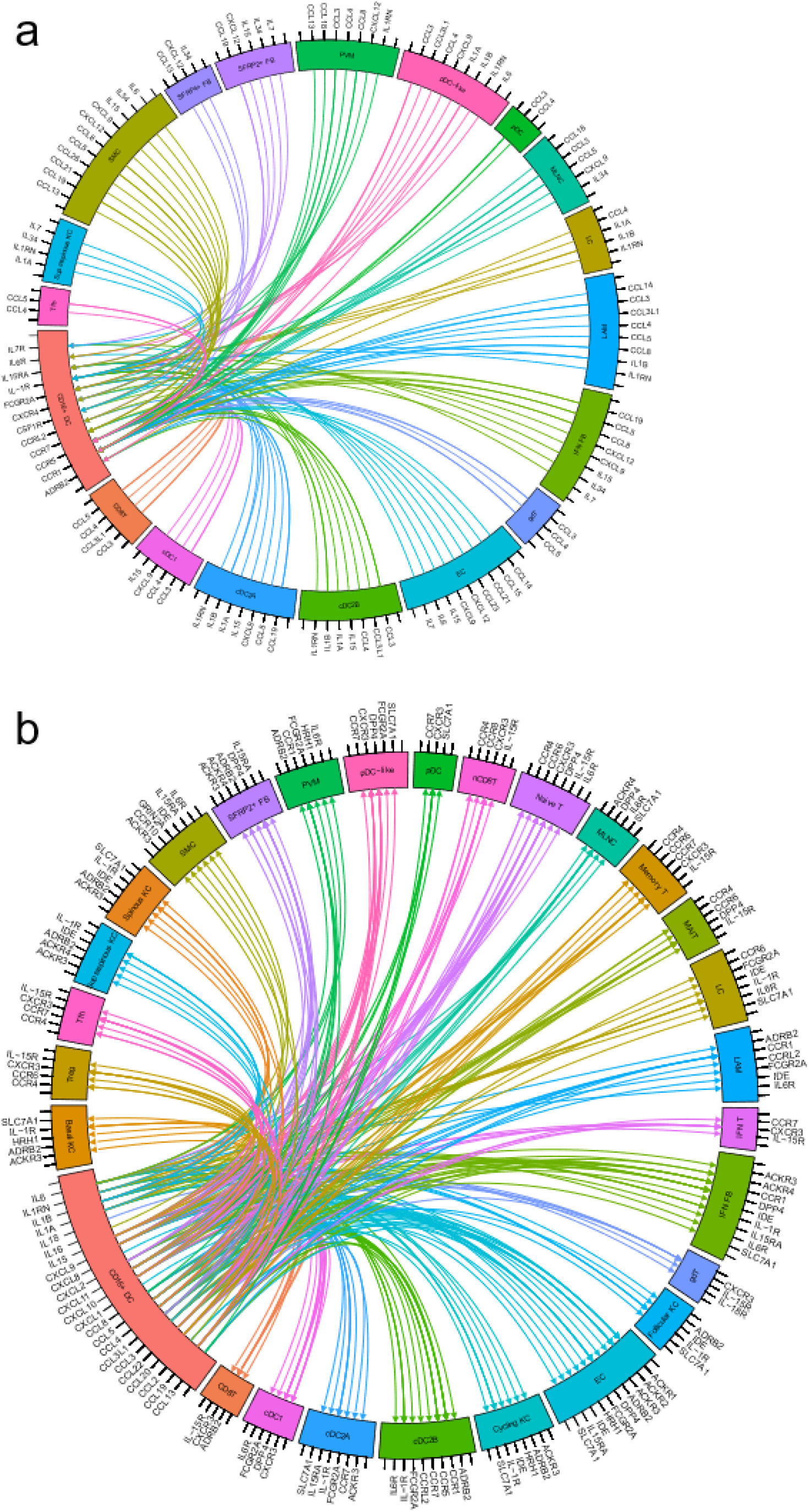
Circos plots showing ligand-receptor (L-R) pairs involving CD16+ DCs, divided by cell subset. Arrows indicate direction from ligand to receptor. **a.** L-R pairs in which CD16+ DCs express the receptor. **b.** L-R pairs in which CD16+ DCs express the ligand.

## SUPPLEMENTARY TABLES

**Supplementary Table 1.** Cutaneous lupus erythematosus (CLE) patient characteristics. DLE, discoid lupus erythematosus. SCLE, subacute cutaneous lupus erythematosus. ITP, immune thrombocytopenia. APS, antiphospholipid syndrome.

| Age (y) | Sex | Active disease manifestations | All current and prior disease manifestations | Years since lupus diagnosis |
| --- | --- | --- | --- | --- |
| 42 | Female | Skin rash (DLE), ITP | Retinal vasculitis, thrombotic vasculopathy, arthritis, ITP, skin rash | 15 |
| 42 | Female | Skin rash (DLE) | Cerebritis (confusion, amnesia), Class IV nephritis, vasculitis, oral ulcers, APS, skin | 20 |
| 48 | Female | Skin rash (SCLE) | Arthritis, sicca symptoms, skin rash, Class V nephritis, serositis | 30 |
| 48 | Female | Skin rash (SCLE), arthritis | Photosensitive rash, Raynaud's phenomenon, arthritis | 4 |
| 60 | Female | Skin rash (DLE) | Photosensitive rash, oral ulcers | 5 |
| 64 | Male | Skin rash (SCLE) | Skin rash | 3 |
| 65 | Female | Photosensitive skin rash | Photosensitive rash, oral ulcers, Raynaud's phenomenon, APS | 1<br>(APS diagnosis 5) |

**Supplementary Table 2. IFN fibroblast (FB) gene markers.** p_val, unadjusted p value; avg_logFC, average log fold change in expression of the indicated gene in IFN FB vs. all other FBs; pct.1, percent of IFN FBs expressing the indicated gene; pct.2, percent of all other FBs expressing the indicated gene; p_val_adj, Benjamini-Hochberg-adjusted p value. Only gene markers with adjusted p values <.05 are included.

**Supplementary Table 3.**
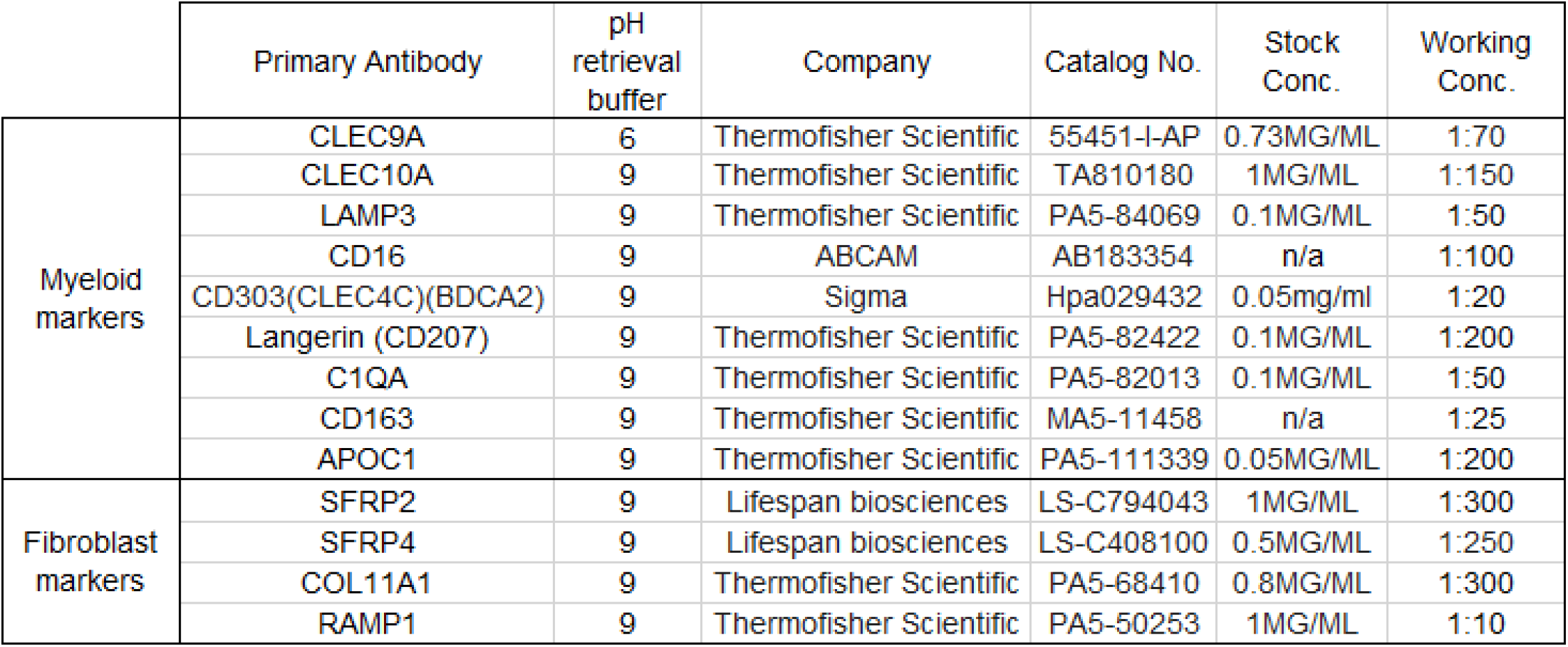
Antibodies used for immunohistochemistry.

**Supplementary Table 4.**
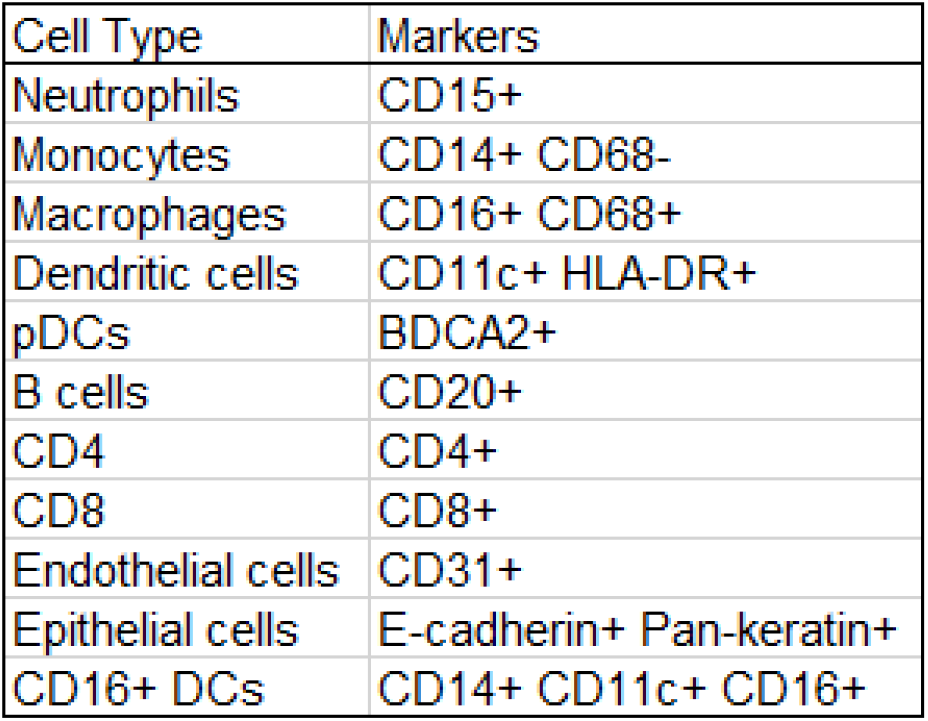
Cell type annotation for imaging mass cytometry.

| Cell Type | Markers |
| --- | --- |
| Neutrophils | CD15+ |
| Monocytes | CD14+ CD68- |
| Macrophages | CD16+ CD68+ |
| Dendritic cells | CD11c+ HLA-DR+ |
| pDCs | BDCA2+ |
| B cells | CD20+ |
| CD4 | CD4+ |
| CD8 | CD8+ |
| Endothelial cells | CD31+ |
| Epithelial cells | E-cadherin+ Pan-keratin+ |
| CD16+ DCs | CD14+ CD11c+ CD16+ |

## REFERENCES

1. Chasset, F. et al. The effect of increasing the dose of hydroxychloroquine (HCQ) in patients with refractory cutaneous lupus erythematosus (CLE): An open-label prospective pilot study. Journal of the American Academy of Dermatology (2016).

2. Sarkar, M.K. et al. Photosensitivity and type I IFN responses in cutaneous lupus are driven by epidermal-derived interferon kappa. Ann Rheum Dis 77, 1653–1664 (2018).

3. Stannard, J.N. et al. Lupus Skin Is Primed for IL-6 Inflammatory Responses through a Keratinocyte-Mediated Autocrine Type I Interferon Loop. The Journal of investigative dermatology 137, 115–122 (2017).

4. Psarras, A. et al. Functionally impaired plasmacytoid dendritic cells and non-haematopoietic sources of type I interferon characterize human autoimmunity. Nat Commun 11, 6149 (2020).

5. Der, E. et al. Tubular cell and keratinocyte single-cell transcriptomics applied to lupus nephritis reveal type I IFN and fibrosis relevant pathways. Nat Immunol 20, 915–927 (2019).

6. Shipman, W.D. et al. A protective Langerhans cell-keratinocyte axis that is dysfunctional in photosensitivity. Science translational medicine 10 (2018).

7. Tabib, T., Morse, C., Wang, T., Chen, W. & Lafyatis, R. SFRP2/DPP4 and FMO1/LSP1 Define Major Fibroblast Populations in Human Skin. The Journal of investigative dermatology 138, 802–810 (2018).

8. Valencia, X., Yarboro, C., Illei, G. & Lipsky, P.E. Deficient CD4+CD25high T regulatory cell function in patients with active systemic lupus erythematosus. J Immunol 178, 2579–2588 (2007).

9. Bonelli, M. et al. Quantitative and qualitative deficiencies of regulatory T cells in patients with systemic lupus erythematosus (SLE). Int Immunol 20, 861–868 (2008).

10. Vargas-Rojas, M.I., Crispin, J.C., Richaud-Patin, Y. & Alcocer-Varela, J. Quantitative and qualitative normal regulatory T cells are not capable of inducing suppression in SLE patients due to T-cell resistance. Lupus 17, 289–294 (2008).

11. Lin, S.C. et al. The quantitative analysis of peripheral blood FOXP3-expressing T cells in systemic lupus erythematosus and rheumatoid arthritis patients. Eur J Clin Invest 37, 987–996 (2007).

12. Kalekar, L.A. & Rosenblum, M.D. Regulatory T cells in inflammatory skin disease: from mice to humans. Int Immunol 31, 457–463 (2019).

13. Wolf, S.J. et al. Ultraviolet light induces increased T cell activation in lupus-prone mice via type I IFN-dependent inhibition of T regulatory cells. J Autoimmun 103, 102291 (2019).

14. Yan, B. et al. Dysfunctional CD4+,CD25+ regulatory T cells in untreated active systemic lupus erythematosus secondary to interferon-alpha-producing antigen-presenting cells. Arthritis Rheum 58, 801–812 (2008).

15. Schiffer, L., Worthmann, K., Haller, H. & Schiffer, M. CXCL13 as a new biomarker of systemic lupus erythematosus and lupus nephritis - from bench to bedside? Clin Exp Immunol 179, 85–89 (2015).

16. Arazi, A. et al. The immune cell landscape in kidneys of patients with lupus nephritis. Nat Immunol 20, 902–914 (2019).

17. Rao, D.A., Arazi, A., Wofsy, D. & Diamond, B. Design and application of single-cell RNA sequencing to study kidney immune cells in lupus nephritis. Nat Rev Nephrol 16, 238–250 (2020).

18. Villani, A.C. et al. Single-cell RNA-seq reveals new types of human blood dendritic cells, monocytes, and progenitors. Science 356 (2017).

19. Bos, J.D., van Garderen, I.D., Krieg, S.R. & Poulter, L.W. Different in situ distribution patterns of dendritic cells having Langerhans (T6+) and interdigitating (RFD1+) cell immunophenotype in psoriasis, atopic dermatitis, and other inflammatory dermatoses. The Journal of investigative dermatology 87, 358–361 (1986).

20. Sontheimer, R.D. & Bergstresser, P.R. Epidermal Langerhans cell involvement in cutaneous lupus erythematosus. The Journal of investigative dermatology 79, 237–243 (1982).

21. MacDonald, K.P. et al. Characterization of human blood dendritic cell subsets. Blood 100, 4512–4520 (2002).

22. Piccioli, D. et al. Functional specialization of human circulating CD16 and CD1c myeloid dendritic-cell subsets. Blood 109, 5371–5379 (2007).

23. Henriques, A. et al. Functional characterization of peripheral blood dendritic cells and monocytes in systemic lupus erythematosus. Rheumatol Int 32, 863–869 (2012).

24. Rhodes, J.W., Tong, O., Harman, A.N. & Turville, S.G. Human Dendritic Cell Subsets, Ontogeny, and Impact on HIV Infection. Front Immunol 10, 1088 (2019).

25. Hansel, A. et al. Human 6-sulfo LacNAc (slan) dendritic cells have molecular and functional features of an important pro-inflammatory cell type in lupus erythematosus. J Autoimmun 40, 1–8 (2013).

26. Dobel, T. et al. FcgammaRIII (CD16) equips immature 6-sulfo LacNAc-expressing dendritic cells (slanDCs) with a unique capacity to handle IgG-complexed antigens. Blood 121, 3609–3618 (2013).

27. Liu, J., Berthier, C.C. & Kahlenberg, J.M. Enhanced Inflammasome Activity in Systemic Lupus Erythematosus Is Mediated via Type I Interferon-Induced Up-Regulation of Interferon Regulatory Factor 1. Arthritis Rheumatol 69, 1840–1849 (2017).

28. Calzetti, F. et al. Human dendritic cell subset 4 (DC4) correlates to a subset of CD14(dim/−)CD16(++) monocytes. J Allergy Clin Immunol 141, 2276–2279 e2273 (2018).

29. Dutertre, C.A. et al. Single-Cell Analysis of Human Mononuclear Phagocytes Reveals Subset-Defining Markers and Identifies Circulating Inflammatory Dendritic Cells. Immunity 51, 573–589 e578 (2019).

30. Aringer, M. et al. 2019 European League Against Rheumatism/American College of Rheumatology classification criteria for systemic lupus erythematosus. Annals of the Rheumatic Diseases 78, 1151–1159 (2019).

31. Butler, A., Hoffman, P., Smibert, P., Papalexi, E. & Satija, R. Integrating single-cell transcriptomic data across different conditions, technologies, and species. Nat Biotechnol 36, 411–420 (2018).

32. Korsunsky, I. et al. Fast, sensitive and accurate integration of single-cell data with Harmony. Nat Methods 16, 1289–1296 (2019).

33. Gudjonsson, J.E. et al. Contribution of plasma cells and B cells to hidradenitis suppurativa pathogenesis. JCI Insight 5 (2020).

34. Efremova, M., Vento-Tormo, M., Teichmann, S.A. & Vento-Tormo, R. CellPhoneDB: inferring cell-cell communication from combined expression of multi-subunit ligand-receptor complexes. Nat Protoc 15, 1484–1506 (2020).

35. Trapnell, C. et al. The dynamics and regulators of cell fate decisions are revealed by pseudotemporal ordering of single cells. Nat Biotechnol 32, 381–386 (2014).

36. Schapiro, D. et al. histoCAT: analysis of cell phenotypes and interactions in multiplex image cytometry data. Nat Methods 14, 873–876 (2017).

